# MIA-Jet: Multi-scale Identification Algorithm of Chromatin Jets

**DOI:** 10.1101/2025.08.27.672730

**Authors:** Sion Kim, Minji Kim

## Abstract

The mammalian genome is organized into large-scale chromosome territories, compartments, domains, and at the smallest scale, chromatin loops and stripes. The newest element is a chromatin jet, a diffused line perpendicular to the main diagonal in the Hi-C contact map, which was reported in quiescent mammalian lymphocytes supporting a two-sided symmetric cohesin loop extrusion model. A similar structure is observed in Repli-HiC and related data, where relatively thin and straight chromatin fountains indicate coupling of DNA replication forks. However, the precise biological implications of these jet-like structures are unknown due to the limitations in computational methods. We developed MIA-Jet, a multi-scale ridge detection algorithm that can accurately detect jets of variable lengths, widths, and angles. When tested on Hi-C, Repli-HiC, ChIA-PET, ChIA-Drop, and Micro-C data in mouse, human, roundworm, and zebrafish cells, MIA-Jet outperformed existing methods. In human cells, jets were enriched in cohesin loading sites and early replication initiation zones. Applying MIA-Jet to Hi-C data generated from protein-degraded cells revealed that jets are dependent on cohesin and that depleting CTCF results in longer and less angled jets than wild-type. We envision MIA-Jet to be broadly applicable to any 3D genome mapping data, thereby providing new insights into the functional roles of chromatin jets.

## INTRODUCTION

The advancement in high-throughput three-dimensional (3D) genome mapping technologies revealed that the mammalian genome is highly organized within the 3D nuclear space. At the largest scale, chromosomes occupy distinct territories, which encompass compartments and topologically associating domains (Lieberman-Aiden et al., 2009; Dixon et al., 2012) (**Figure 1a**, ‘TAD’). At a smaller scale, various protein factors mediate chromatin loops (**Figure 1a**, ‘Loop’) to bridge distal genomic loci together (Fullwood et al., 2009; Li et al., 2012; Tang et al., 2015; Weintraub et al., 2017; Mumbach et al., 2016; Grubert et al., 2020) where loops are sometimes accompanied by architectural stripes (Vian et al., 2018) (**Figure 1a**, ‘Stripe’). Each of these chromatin structures has potential roles in cellular processes. In particular, chromatin loops are considered the basic unit of higher order genome organization and are extensively studied (Phillips-Cremins et al., 2013; Braccioli and de Wit, 2019; Davidson et al., 2023). For instance, more than 22 computational tools have helped to annotate and characterize chromatin loops in 3C data (Chowdhury et al., 2024). It is commonly known that cohesin extrudes DNA until it is blocked by a zinc finger protein CTCF, which binds to specific DNA motifs. Most of the chromatin loops are anchored at a pair of convergent CTCF binding motifs (Rao et al., 2014; Tang et al., 2015), suggesting that the orientation of binding motifs is related to the biophysical mechanism of chromatin loop formation. Notably, perturbing CTCF sites alters genome organization and has implications in cancer and genetic disorders (Ushiki et al., 2021; Kubo et al., 2021), highlighting the importance of chromatin loops in maintaining the normal function of cells.

**Figure 1:**
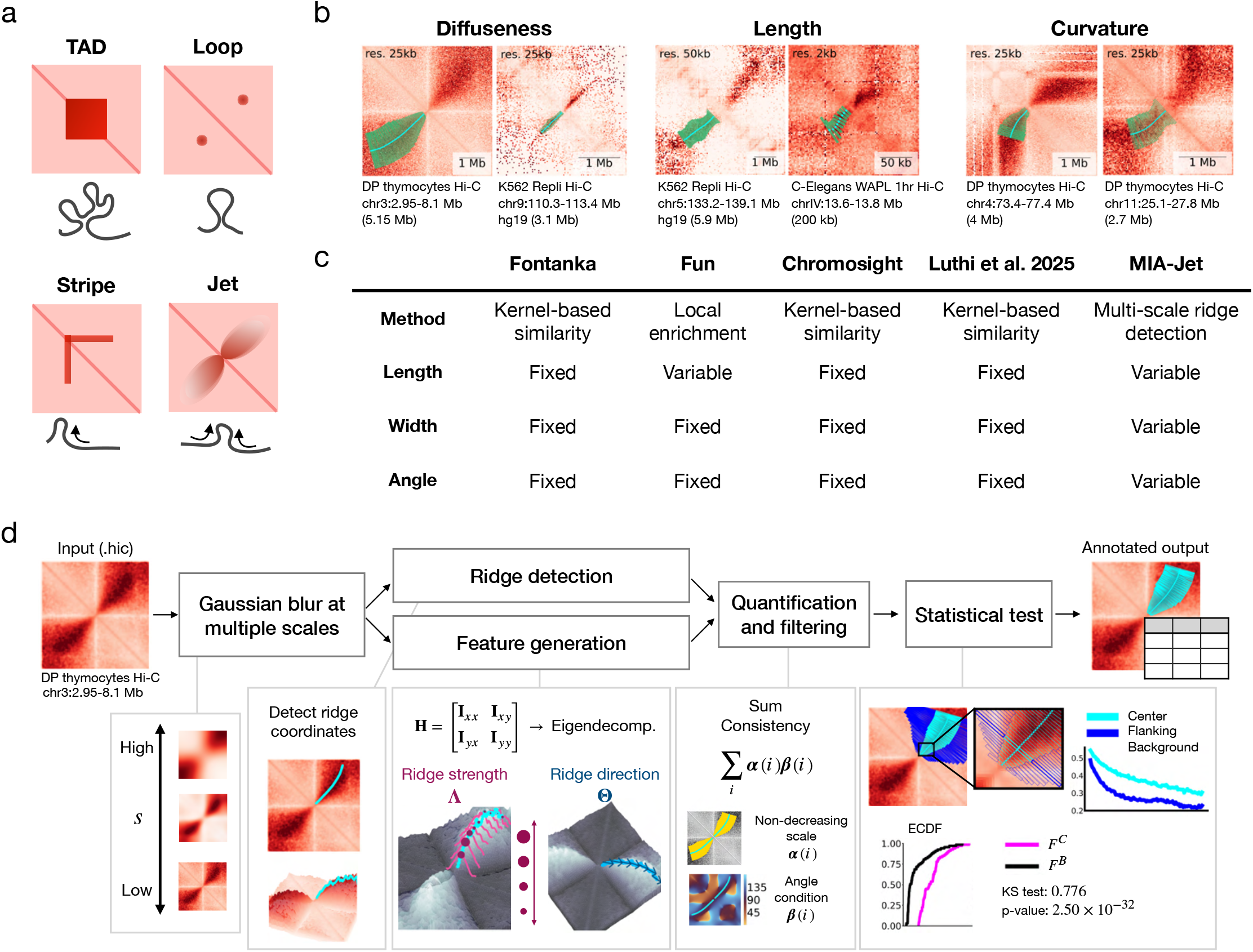
Overview of MIA-Jet algorithm. **(a)** Schematic 2D contact map showing TAD, loop, stripe, and jet. The corresponding chromatin configuration for each structure is shown below. **(b)** Observed/expected 2D contact maps with MIA-Jet output annotation in the lower triangle: jet coordinates (cyan line) and jet widths (green boxes). **(c)** Summary table of available jet calling algorithms Fontanka, Fun, Chromosight, Luthi et al., 2025, and MIA-Jet. **(d)** MIA-Jet pipeline detailing the major components of the algorithm. The input .hic file undergoes Gaussian blurring at multiple scales. The blurred Hi-C maps are then passed into two modules: (1) ridge detection, which infers the coordinates of the jets, and (2) feature generation, where eigen-decomposition of the image Hessian matrix H is used to obtain ridge strength *Λ* and ridge direction **Θ**. Features are combined with coordinates obtained from ridge detection to rank each ridge by “sum consistency”, a weighted sum of softly imposed constraints, such as a specific width profile *α* and the desired angle of jet extrusion **β**. Finally, a statistical test is performed on a local enrichment ratio compared to the flanking regions. The output is a table with jet position, angle, width and quantification metrics. See also **Figure S1**.

Understanding the fundamental biophysical principle of chromatin loop formation is critical for cataloging their functional roles in cellular processes such as gene transcription. However, the extrusion mechanism remains largely unsettled. Cohesin was proposed to extrude DNA loops symmetrically in a two-sided manner *in vitro* through biochemical and single-molecule imaging methods (Davidson et al., 2019; Kim et al., 2019), yet a recent *in vitro* study supported an asymmetric one-sided extrusion model instead (Barth et al., 2025). The problem is equally perplexing *in vivo*. Architectural stripes (Vian et al., 2018) imply that cohesin extrusion occurs asymmetrically at one side of the CTCF loop anchoring sites, while chromatin jets (**Figure 1a**, ‘Jet’) reported in Hi-C data of quiescent mouse cells (Guo et al., 2022) and *in silico* simulations (Banigan et al., 2020) posit that cohesin extrudes loops in a two-sided symmetric fashion upon being loaded onto the DNA by loading factors including NIPBL. These isolated results have raised debates regarding the model of cohesin extrusion *in vivo* (Drayton and Hansen, 2022). A key question is whether chromatin jets are widely observed in mammalian cells or are specific to quiescent mouse cells, but it is unclear as the 37 jets were manually annotated without a computational algorithm to automatically detect jets.

Subsequent studies also revealed jet-like patterns—sometimes referred to as fountains or plumes—in multiple species using various 3D genome mapping technologies. In *C. elegans*, chromatin fountains at active enhancer sites disappear upon cohesin cleavage (Luthi et al., 2025). Another study using auxin-inducible degron system confirmed that jets are dependent on cohesin and that they emerge from NIPBL binding (cohesin loading) sites; depleting a cohesin unloading factor WAPL in *C. elegans* resulted in the extension of jets (Kim et al., 2025), akin to the extended loops observed in human cells (Haarhuis et al., 2017). Performing Hi-C experiments in zebrafish embryos showed that fountains emerge selectively at enhancers upon zygotic genome activation and the authors attribute the findings to two-sided, desynchronized loop extrusion by cohesin upon loading onto the DNA (Galitsyna et al., 2026). Jets have also been reported in fungi (Shao et al., 2024) and mouse embryonic stem cells (Liu et al., 2025).

The aforementioned results are derived from Hi-C data analysis and unanimously suggest that jets are: 1) found in various species, 2) often associated with active enhancer or cohesin loading sites, and 3) dependent on cohesin. In parallel, a new replication-associated in situ Hi-C method (Repli-HiC) was developed to enrich for chromatin interactions involving nascent DNA (Liu et al., 2024), similar to ChIA-PET (Fullwood et al., 2009), HiChIP (Mumbach et al., 2016), and PLAC-seq (Fang et al., 2016) that enrich for protein-mediated interactions. By applying Repli-HiC to human cells, the authors identified two classes of chromatin fountains: Type I fountains (median length of 160 kb) favoured at early replication initiation zones, and Type II fountains (median length of 1 Mb) associated with replication termination (Liu et al., 2024). More recently, Repli-MiC (Zhangding et al., 2026) improved the resolution of Repli-HiC method by incorporating the micrococcal nuclease digestion, while Rep-Hi-C (Gaggioli et al., 2026) has been applied to various human cell lines. Collectively, these results provide convincing evidence that the chromatin jets observed by Guo et al. in 2022 may be a genome-wide phenomena across various technologies and species, although the extent to which jets are due to cohesin extrusion or DNA replication remains unknown.

Current progress is impeded by a lack of computational tools to identify and quantify jets in an unbiased, genome-wide manner. The 37 jets in mouse cells were manually annotated by hand (Guo et al., 2022), while Repli-HiC fountains were identified by a local enrichment method called Fun (Liu et al., 2024); Repli-MiC method was accompanied by a reinforcement learning approach Fun2 (Zhangding et al., 2026). Other studies have computationally identified jets by kernel-based similarity methods Fontanka (Galitsyna et al., 2026), Chromosight (Matthey-Doret et al., 2020), and Luthi et al., 2025. These methods require well-defined kernels, which may require hand annotation by users. The method used by Luthi et al., 2025 also mandates cohesin-depleted Hi-C data as an input to calculate the expected background distribution, which may not be widely available in cell lines due to technical difficulties and cost. Most importantly, current methods are limited in identifying jets of fixed width and angle, thereby posing a computational challenge in finding diffused or curved jets reported in Guo et al., 2022. Kernel-based methods also assume fixed length, which is not suitable for Repli-HiC fountains as Type I and Type II fountains differ in length (median of 160 kb vs. 1 Mb; Liu et al., 2024). Therefore, existing computational methods cannot be broadly applied to identify jets of various forms. As a result, it is challenging to answer important biological questions, such as: if jets are found in all cells, species, across technologies; whether jets coincide with cohesin loading sites for loop extrusion and/or DNA replication origins; if depleting protein factors result in changes in the number, strength, angle, and length of jets.

To overcome the technical limitations of existing algorithms and to answer biological questions, we developed MIA-Jet, a multi-scale ridge detection algorithm to accurately identify jets of variable length, width, and angle. It is inspired by the earlier success of computer vision-based methods MUSTACHE (Roayaei Ardakany et al., 2020), SIP (Rowley et al., 2020), and Stripenn (Yoon et al., 2022) to call chromatin loops or architectural stripes. When tested on simulated data, Hi-C data in quiescent mouse cells, and Repli-HiC data in human K562 cells, MIA-Jet outperformed existing algorithms Fun and Fontanka with respect to known jet annotations and genomic features such as NIPBL binding intensity. MIA-Jet identified jets across various technologies—in situ Hi-C, intact Hi-C, Repli-HiC, ChIA-PET, ChIA-Drop, Micro-C—in human, mouse, *C. elegans*, and zebrafish (**Supplementary Table S1**; Reiff et al., 2022; Luo et al., 2020; Rao et al., 2017; Rao et al., 2014; Liu et al., 2024; Kieffer-Kwon et al., 2017; Guo et al., 2022; Hsieh et al., 2022; Kim et al., 2025; Morao et al., 2022; Wike et al., 2021).

Compared to the random background locations, jets identified by MIA-Jet had high insulation scores, were enriched in NIPBL binding signal and enhancer marks, and were at or near early DNA replication initiation zones. Finally, by using publicly available Hi-C and Micro-C datasets derived from protein-depleted cells, we show that depleting cohesin eliminates jets, while co-degrading CTCF and WAPL in mouse embryonic stem cells results in stronger and longer jets; CTCF depletion yields longer and less angled jets. We envision MIA-Jet to be instrumental in answering key biological questions in the 3D genomics field as it is versatile and applicable to accommodate a wide range of genome mapping technologies, species, and quantitative features of jets.

## RESULTS

### Overview of the MIA-Jet algorithm

Chromatin jets pose unique computational challenges that are distinct from other chromatin features such as TADs, loops, and stripes, for which there already exist a plethora of available callers (Xu et al., 2024; Chowdhury et al., 2024; Raffo and Paulsen 2023). First, jets can have different width profiles, such as “diffuse” jets observed in DP thymocyte Hi-C data where the signal gradually decays with genomic distance, or those that exhibit sharp, edge-like boundaries in the case of K562 Repli-HiC data (**Figure 1b**, ‘Diffuseness’). Second, jets vary in length: some jets are as large as 4 Mb while others are small in the order of 50 kb (**Figure 1b**, ‘Length’).

Third, unlike architectural stripes that are either horizontal or vertical, jets may extrude at variable angles and exhibit curvature as the jet protrudes (**Figure 1b**, ‘Curvature’). Existing computational tools cannot accommodate all three features of jets. Fun (Liu et al., 2024) scans through the genome and computes local enrichment of jet-like structures to identify Type I and Type II fountains in Repli-HiC data by allowing users to specify the expected length of the fountains but yields jets of fixed width and angle (**Figure 1c**). This method was improved upon with Fun2 (Zhangding et al., 2026), which uses a reinforcement learning framework to allow variable width and angled jets. However, the model only associates a jet with a single angle parameter, lacking the ability to capture jets that have different curvatures at various points in the extrusion process. Fontanka (Galitsyna et al., 2026), Chromosight (Matthey-Doret et al., 2020), and Luthi et al., 2025 requires users to provide the mask, which is either averaged hand-annotated regions or a binary mask, to compute the similarity between the kernel and sliding window across the main diagonal. As these methods rely heavily on the kernel, they can only identify specific types of jets and potentially miss others that deviate from the pre-defined kernel.

Here, we present MIA-Jet, a multi-scale ridge detection algorithm that can accurately identify jets of variable lengths, widths, and angles. We leverage concepts from image processing to define chromatin jets as *ridges*, which is a line on the image surface where the signal is relatively stable along the line but decays to the left and the right on either side (Haralick, 1983). Utilizing local second order derivatives, we bypass the need for users to define kernels that impose one particular shape and size of jet. Ridge detection is combined with scale-space for multi-scale ridge detection and automatic width inference (Lindeberg, 1998). MIA-Jet first extracts a rectangular box along the main diagonal (**Figure S1**, ‘Hi-C image generation’) and applies Gaussian blur at multiple scales to the input Hi-C matrix to generate a stack of images: low blurring keeps fine structures and high blurring preserves large-scale structures (**Figure 1d**). The algorithm then detects ridge coordinates by the *CurveTrace* ImageJ plugin (Katrukha, 2021) (**Figure 1d, S1**, ‘Ridge detection’). Separately, MIA-Jet computes the image Hessian **H** to quantify the ridge strength **Λ** and ridge angle **Θ** (**Figure 1d, S1**, ‘Feature generation’). We then use automatic scale selection (Lindeberg, 1998) to infer the appropriate width of each jet and find the corresponding jet strengths and angles (**Figure S1**, ‘Scale inference’). These metrics are then used to process and rank ridges based on user-defined features, such as the expected width profiles or the desired angle of jet extrusion (**Figure 1d, S1**, ‘Quantification and filtering’). Finally, the algorithm tests for significant enrichment of the observed ridge compared to its local neighborhood and outputs annotated jets with genomic coordinates, corresponding widths, and angles (**Figure 1d**, ‘Annotated output’; **Figure 1b**, lower triangle of 2D contact maps). The MIA-Jet algorithm is implemented into a publicly available python program. Algorithmic details are provided in the **Methods** section.

### Evaluating performance in simulated data highlights MIA-Jet’s accuracy and algorithmic flexibility

We first sought to evaluate the performance of MIA-Jet and existing computational jet callers in an unbiased manner via polymer simulations. Simulated data provides full control of the jet features such as lengths and angles, while also retaining the ground truth jet locations to quantify the performance of algorithms. We generated simulated jets using 3DPolyS-LE (Gitchev et al., 2022) (**Methods**, *Generating simulated data*). Following Guo et al., 2022, where the authors showed that jet-like patterns in Hi-C maps could arise due to the two-sided loop extrusion mechanism by cohesin, we used cohesin loading sites to arbitrarily determine the jet origins and optionally introduced jet curvature by placing CTCF boundary elements at various locations away from the loading site (**Figures 2a, S2a**, row 1).

**Figure 2:**
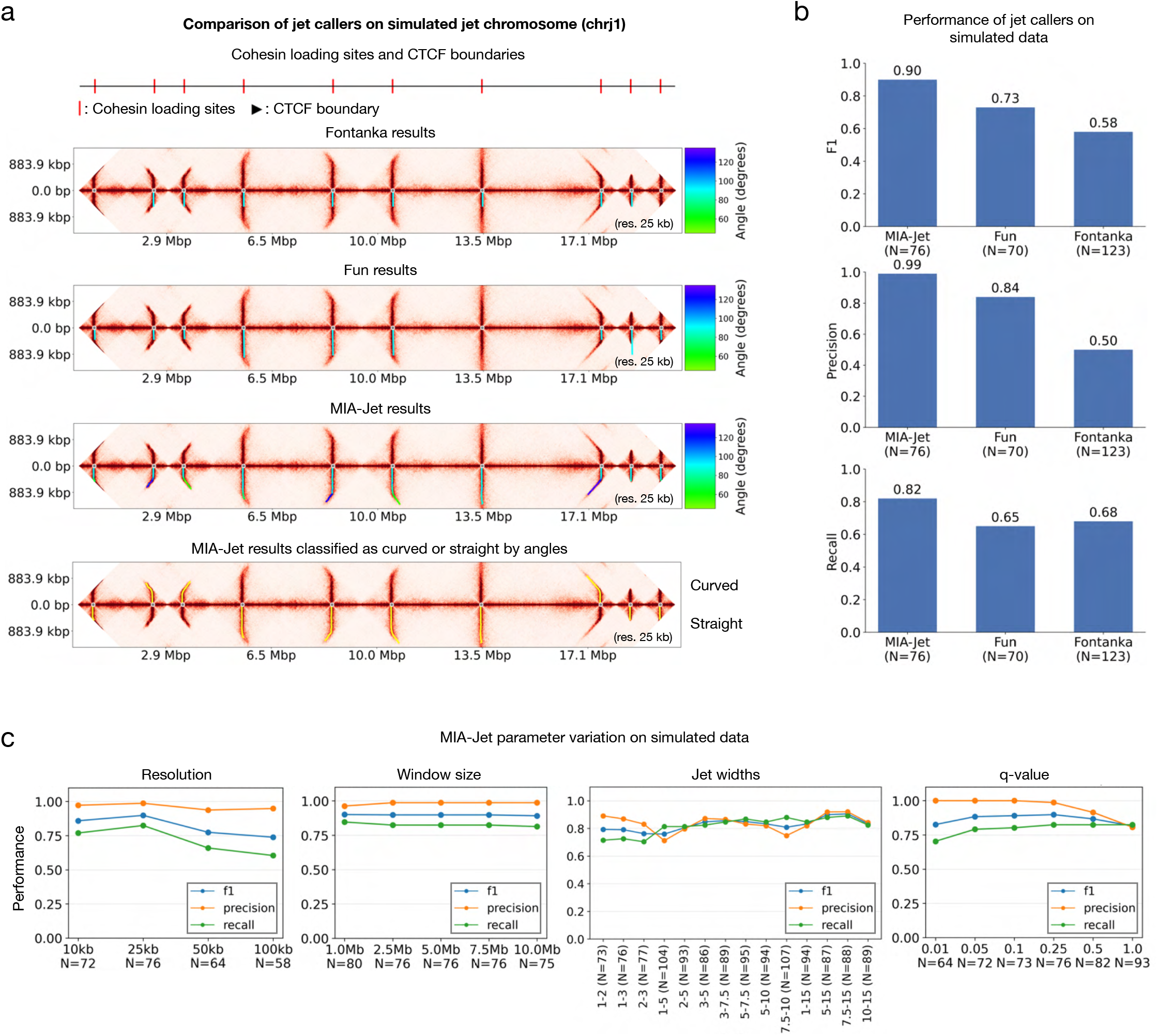
Performance of jet callers on simulated data. **(a)** Example of simulated chromosome chrj1 containing jets (‘test1a_fast’ **Methods**, *Generating simulated data*). Location of ground truth jet loading sites (row 1). Jets called by Fontanka (cyan line) (row 2), Fun (cyan line) (row 3), and MIA-Jet (coloured by angle) (row 4). Jets called by MIA-Jet classified into “Curved” (upper half) or “Straight” (lower half) using angle annotations (row 5). **(b)** Performance of MIA-Jet, Fun, and Fontanka on full simulated dataset. N is the number of jets called by each method. **(c)** Performance of MIA-Jet across parameter variation on simulated data. Jet widths parameter (plot 3) takes a lower bound and upper bound for the expected jet widths (in pixels) (e.g., 1-2 should find jet whose widths are between 1 and 2 pixels wide).

We ran Fontanka, Fun and MIA-Jet and showed their respective annotations in the lower half of each row (**Figure 2a**, row 2-4). Fontanka was selected among the kernel-based methods (**Figure 1c**) as the authors had previously demonstrated that their results are comparable with those of Chromosight (Galitsyna et al., 2026). Qualitative inspection reveals all three methods performed reasonably well in identifying straight, fixed-length jets, but Fun and Fontanka missed jets of variable lengths and curvatures. Notably, Fontanka missed two jets on each ends of the simulated jet chromosome (chrj1) and its annotations are all of the same length (**Figure 2a**, row 2), highlighting the limitations of a fixed size kernel convolution (**Figure 1c**). Comparatively, Fun can dynamically adjust the length annotations based on the size of each jet but failed to identify a long, straight jet near 13.5 Mbp artificial coordinate (**Figure 2a**, row 3). Note that Fontanka and Fun annotated only straight jets of 90 degrees in cyan color (**Figure 2a**, rows 2, 3). MIA-Jet has a unique ability to both track the jet line as it changes direction and quantify the angle at each point, where jets with curvature have angles around 120 degrees as indicated in blue (**Figure 2a**, row 4). Users may utilize these annotations downstream, such as in categorizing jets as “curved” or “straight” based on the range of angles (**Figure 2a**, row 5).

In total, we generated 7 chromosomes collectively containing 91 jets and 2 chromosomes containing TADs as a negative control, as exemplified in **Figure S2a** and **Figure S2b**, respectively. Notably, MIA-Jet correctly identified all jets in chrj2—straight, angled, long, and short—while keeping TADs largely unannotated in chrt1; Fontanka and Fun failed to annotate curved jets or wrongfully reported TADs or non-loading sites as jets (**Figure S2a, S2b**). Beyond these examples, the performance pooled across all simulated data show MIA-Jet had the highest F1 score at 0.90 compared to Fun at 0.73 or Fontanka 0.58 (**Figure 2b**).

Precision and recall plots reveal that MIA-Jet had almost perfect precision (0.99) and comparatively lower recall (0.82), suggesting that MIA-Jet is better at filtering out false jets than finding all the true jets. Nevertheless, MIA-Jet had higher performance metrics in all the categories than Fun or Fontanka (**Figure S2c**). On simulated data at 25kb resolution, MIA-Jet had longer run-time and higher memory usage than Fun and Fontanka, yet all methods ran relatively quickly at less than 5 minutes run-time and 5 GB peak memory consumption (**Figure S2d**). We envision the runtime and memory of MIA-Jet to be reasonable for typical genomics research groups.

We then tested the robustness of MIA-Jet by varying key parameters. Specifically, we varied the resolution from 10kb to 100kb, window size from 1 Mb to 10 Mb, jet widths from 1-2 to 10-15 (lower bound and upper bound in pixels), and q-value threshold from 0.01 to 1 (**Methods**). MIA-Jet had consistent F1 score of around 0.8 or higher throughout all parameter variations (**Figure 2c**). The resolution refers to the bin size of the Hi-C contact map used by the program, the window size is the maximum genomic extent from the main diagonal to capture jets, and jet widths is the desired range (upper and lower bounds) in number of pixels for the jet widths (**Methods**, *Choosing the scales*). Pixel-counting on the Hi-C image suggested a number between 5-10 pixels as the true jet width. As expected, the further the jet widths deviate from 5-10 range, such as at 1-2 or 1-3, we find an impact on performance compared to the other ranges, such as 5-15 (**Figure 2c**). Remarkably, even if the specified jet width range did not overlap or include the true jet width range, the performance of MIA-Jet was relatively consistent with an F1 score of around 0.8 and outperformed Fun (0.73) or Fontanka (0.58) (**Figure 2b, Figure 2c**). Altogether, experiments on simulated data suggest MIA-Jet is robust and reproducible across parameter variation.

As a comparison, we also tested the robustness of Fun and Fontanka. Fun performs well when the resolution is 10kb or 25kb, with F1-score of around 0.7 that drastically decreases as the resolution reaches 50kb and 100kb (**Figure S2e**). This is likely a result of the padding width parameter, which controls the effective width of jets, being directly related to the resolution; at the default of 2 pixels, a resolution of 100kb means 500kb wide sampling box, which may be too large (**Figure S2e**). When the extension length, which specifies the maximum extrusion length of the jets, varies from 1 Mb to 4.2 Mb, Fun stabilizes in performance with an F1 score of 0.7 (**Figure S2e**). Fontanka – despite having lower F1-scores than Fun and MIA-Jet – has consistent performance across resolutions from 10kb to 100kb but is sensitive to varying the window size (**Figure S2f**). For Fontanka, the expected length of jets is directly related to the window size parameter, which is the kernel size of the jet mask and is the likely reason for the sensitivity towards window size.

Overall, MIA-Jet shows superior performance and robustness on simulated data compared to existing jet callers, offering additional features beyond the approximate jet location, such as line and angle annotations.

### MIA-Jet outperforms existing methods on quiescent mouse cells Hi-C data

We ran MIA-Jet, Fun and Fontanka on DP thymocytes Hi-C data, where 37 jets were reported by Guo et al. 2022. A representative jet called by MIA-Jet is shown with line and width annotation (**Figure 3a**). The top 5 jets ranked by ‘sum consistency’ in MIA-Jet include known Guo et al. jets as ranks 2 and 4 (**Figure 3b**, top panel). Top 5 jets ranked by ‘SoN’ (signal over noise) in Fun reported a mixture of jets, stripes and loops (**Figure 3b**, middle panel) whereas Fontanka ranked by ‘FS_peaks’ mostly reported compartments or domains near unmappable regions (**Figure 3b**, bottom panel). Note that none of the top 5 jets called by Fun and Fontanka have captured known Guo et al. jets. A major advantage of MIA-Jet is its ability to annotate the width of jets. For example, a jet found on chromosome 14 is annotated with narrow widths from the origin and gradually increasing in widths, broadening off diagonal (**Figure 3a**). Genome-wide aggregation of jets identified by each method shows comparable aggregate signal strength across the methods. Noticeably, MIA-Jet and Fontanka have similar aggregate jet profiles with the Guo et al. set, although Fontanka is limited in finding only 39 jets (**Figure 3c**) and reported off-diagonal TAD-like blocks instead of jets (**Figure 3b**, bottom panel). In comparison, Fun called more jets than any other method at 579 jets and show a profile that resembles a stencil-like narrow enrichment, differing from the other methods (**Figure 3c**). While MIA-Jet identified 332 jets, about 9-fold more than Guo et al., it best recapitulated diffuse 37 jets by Guo et al. (**Figures 3c, 3b**). In addition, MIA-Jet had highest binding signals in H3K27ac, RAD21, and NIPBL, aligning with the chromatin features of jets reported in Guo et al. (**Figure 3d**). As NIPBL binding is generally indicative of cohesin loading sites from which jets were shown to originate in quiescent mouse cells, we used it as a proxy for a likelihood of a true jet. When comparing MIA-Jet, Fun, and Fontanka against the reference 37 jets identified by Guo et al., we found MIA-Jet to have a higher number of 32 overlapping jets than Fun (24 jets) and Fontanka (8 jets) and comparable NIPBL binding intensity to that of Guo et al., whereas Fun and Fontanka were markedly lower (**Figure 3e**). We then compared among the computational jet-callers and found MIA-Jet missed 444 and 24 jets called by Fun and Fontanka, respectively (**Figure S3a**, top panel). Nevertheless, these Fun-specific and Fontanka-specific jet origins had lower NIPBL signal enrichment than that of MIA-Jet specific jets (**Figure S3a**, bottom panel), implying that Fun and Fontaka exclusively identified jets that are likely not at cohesin loading sites.

**Figure 3:**
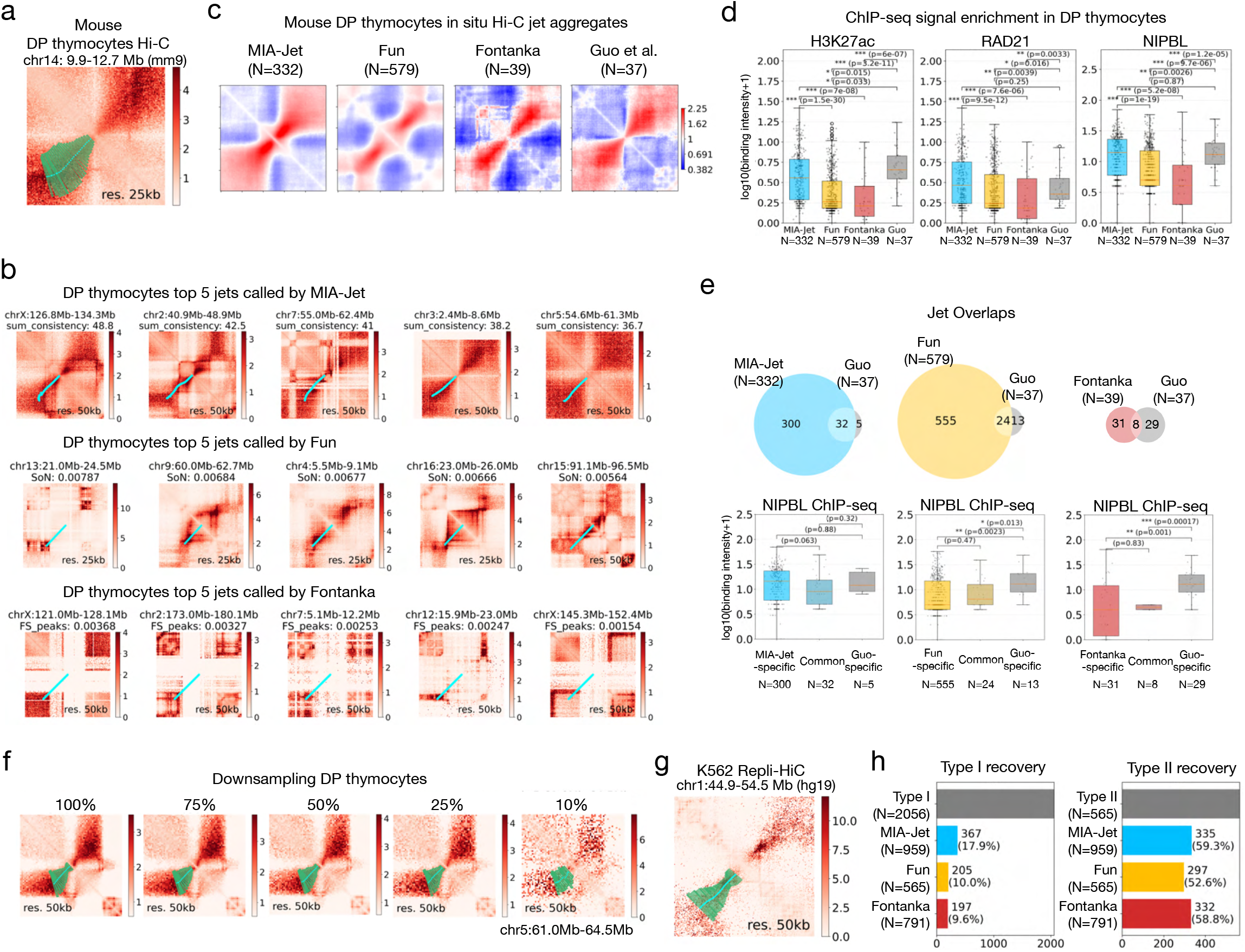
Comparing MIA-Jet, Fun, Fontanka on mouse Hi-C data and human Repli-HiC data. **(a)** Observed/Expected map of a jet in mouse DP thymocyte Hi-C data with MIA-Jet position annotation (cyan line) and width (green boxes). **(b)** Top 5 jets identified by MIA-Jet (row 1), Fun (row 2), and Fontanka (row 3) in DP thymocytes Hi-C data. Maps show Observed/Expected values and annotation in lower triangle. **(c)** Aggregate Observed/Expected maps at jet locations identified by MIA-Jet, Fun, Fontanka, and Guo et al., 2022, in mouse DP thymocytes in situ Hi-C data. **(d)** Boxplots of log-transformed H3K27ac, RAD21, and NIPBL ChIP-seq data in mouse DP thymocytes at jet locations identified by each method (p-values from two-sided Wilcoxon rank-sum test). **(e)** Comparison of jets called by MIA-Jet, Fun, and Fontanka with the reference Guo et al. Venn Diagram shows the number of jets common and specific between two methods. The log-transformed NIPBL ChIP-seq boxplot is shown for the common and specific jets below. **(f)** An example jet across downsampled DP thymocyte data at 75%, 50%, 25%, and 10% of the original number of reads (100%); Observed/Expected map with MIA-Jet position annotation (cyan line) and width (green boxes) in lower triangle. (**g**) An example of K562 Repli-HiC fountain identified by MIA-Jet; Observed/Expected map with MIA-Jet position annotation (cyan line) and width (green boxes) in lower triangle. (**h**) Bar plot showing the number of jets called by MIA-Jet, Fun, or Fontanka overlapping with Type I jets (left) and Type II jets (right).

A major concern with computational 3D genome structure callers is their robustness to sequencing depth, which we next set out to test. We generated downsampled DP thymocytes Hi-C data at 75%, 50%, 25% and 10% of the original number of reads (100%). Visual inspection of one exemplary jet shows that MIA-Jet identified this jet with similar lengths and widths until reaching 10%, where the annotated jet is a bit wider and shorter than those of 100%, 75%, 50%, and 25% (**Figure 3f**). Genome-wide, MIA-Jet identified a similar number of jets (271 to 312) until reaching 10%, when it yielded 512 jets (**Figure S3b**). Fun and Fontanka were consistent across all sequencing depth, ranging from 451 to 588, and 29 to 38, respectively (**Figure S3b**). As a biological validation, we examined NIPBL binding intensity at jet locations across sequencing depth, where MIA-Jet shows higher signal than Fun and Fontanka (**Figure S3c**). Furthermore, this separation is maintained from 100% to 25% sequencing depth, then in 10% sequencing depth drops to a comparable value (0.7) to other methods, suggesting that the sudden increase in jets called in 10% by MIA-Jet are likely false positives. In terms of the consistency of results with respect to various sequencing depth, MIA-Jet in general has lower consistency than the other methods, although all three were most vulnerable at 10% sequencing depth (**Figure S3d**).

Mouse splenic B cell Hi-C data also exhibited a jet on chromosome 3 (**Figure S3e**, left) as showcased by Guo et al. in DP thymocytes (**Figure 1b**, left-most), but the extent to which jets are conserved between the two cell types was unknown due to the lack of computational jet callers. By applying MIA-Jet to B cells, we identified jets at this and other locations, such as slightly curved jet on chromosome 4 (**Figure S3e**, middle). In total, we identified 346 jets with a similar aggregate profile as DP thymocytes (**Figure S3e**, right). We then directly compared DP thymocytes and B cell and found 135 shared jets with both shared and cell-type-specific categories exhibiting strong jet-like patterns (**Figure S3f**). Collectively, these results highlight MIA-Jet’s ability to identify jets with diffuse width profiles present in quiescent mouse cells and recapitulate epigenomic features therein.

### MIA-Jet can identify Type I and Type II fountains in K562 Repli-HiC data

In human K562 cell lines, Liu et al. identified 2056 Type I and 565 Type II jets associated with DNA replication (jets also referred to as *fountains* in Liu et al., 2024). Type I jets originate from replication initiation zones and is generally narrow with straight extrusion perpendicular from the main diagonal, and smaller than Type II jets. Comparatively, Type II jets are associated with replication termination zones and have a larger, wider profile than Type I jets. Notably, both Type I and Type II jets have been identified using Fun, with 5kb, 10kb and 20kb runs constituting Type I and 50kb, 100kb, and 200kb for Type II. Both sets were further filtered for association with DNA replication marks (see **Methods** for descriptions).

We then ran MIA-Jet, Fun, and Fontanka at 50 kb in K562 Repli-HiC data and tested whether they can capture these different types—Types I and II—of jets related to DNA replication. A representative Type II jet called by MIA-Jet is shown (**Figure 3g**). In contrast to the diffuse widths reported in mouse quiescent cells (**Figure 3a**), MIA-Jet accurately captured the sharp boundaries of the jet widths in K562 Repli-HiC data (**Figure 3g**). Top 5 jets called by MIA-Jet were large and wide, resembling Type II jets (**Figure S3g**, top panel). Indeed, all top 5 jets called by MIA-Jet corresponded to Type II Liu et al. annotations, while Fun and Fontanka included false positives in the top rankings. Out of the top 5 jets, Fun identified 3 true jets – ranks 1 (Type II), 3 (Type I) and 5 (Type I) – and included 2 false positives while Fontanka identified only 2 true jets, ranks 3 (Type I and II) and 5 (Type I) (**Figure S3g**). We then performed genome-wide evaluation by how well each method can recall Type I or Type II jets. MIA-Jet can recover the highest proportion of both Type I and Type II jets at 367 (17.9%) and 335 (59.3%), respectively (**Figure 3h**). Remarkably, even though all methods were run at a resolution of 50kb, MIA-Jet found more Type I jets than Type II jets, highlighting its multi-scale capabilities. In contrast, Fun and Fontanka found more Type II jets than Type I jets as expected due to the low resolution at 50 kb emphasizing larger scale structures (297 vs 205, 332 vs. 197, respectively) (**Figure 3h**).

When we aggregated Hi-C contacts over the jets called by each method, all three methods had comparably strong signals, where MIA-Jet identified more jets (N=959) than Fun (N=565) and Fontanka (N=791) (**Figure S3h**). Among the additional 581 jets MIA-Jet identified that Fun did not, the aggregate map reveals an elongated jet-like enrichment, while Fun-specific jets resemble dots rather than jets and are relatively weaker in strength (**Figure S3i**, top). The comparison between MIA-Jet and Fontanka shows similar jet shapes but stronger enrichment for MIA-Jet-specific than Fontanka-specific jets (**Figure S3i**, middle). Comparing Fun and Fontanka shows that both methods are similar in jet strength, with Fun-specific jets being slightly thicker than Fontanka-specific jets (**Figure S3i**, bottom). In terms of run time, MIA-Jet, Fun, and Fontanka ran within 7 minutes on DP thymocyte Hi-C data and 12 minutes on K562 Repli-HiC data (**Figure S3j**). Peak memory consumption was low and all methods ran with less than 10 GB of peak memory consumption (**Figure S3j**).

In combination with the results from simulations, applying MIA-Jet to mouse DP thymocytes Hi-C, mouse splenic B cell Hi-C, and K562 Repli-HiC data suggests the algorithm can efficiently and accurately detect jets of various lengths, angles, and widths, thereby motivating us to utilize MIA-Jet to comprehensively characterize jets across various technologies and cell types to uncover biological features of jets in publicly available datasets that have not been fully analyzed for studying jets.

### Repli-HiC has longer and stronger jets than Hi-C

The Repli-HiC method enriches for chromatin interactions associated with nascent DNA among all interactions present in Hi-C experiments, resulting in a subset of contacts. Thus, in theory, Hi-C experiments should also have jets, which are simply buried among all other chromatin structures, as shown in exemplary locations in Liu et al., 2024. We sought to systematically compare the presence, strengths, and overlaps of jets in Repli-HiC, in situ Hi-C, and intact Hi-C in K562 cells by calling jets via MIA-Jet in an unbiased manner. Since the resolution at which we call jets affects the locational uncertainty of the annotations (e.g., ± 2 bins), we ran MIA-Jet at a higher resolution at 25kb to more precisely analyze genomic signals associated with jets.

As expected, Repli-HiC had 3-fold more jets (N=1002) than in situ Hi-C (N=313) and intact Hi-C (N=275), and had the strongest jet signals (**Figure 4a**, top panel). Repli-HiC jets were also longer (median = 375kb; mean = 532kb) than in situ Hi-C (median = 275kb; mean = 339kb) and intact Hi-C (median=240kb; mean=315kb). A jet found in Repli-HiC data is also recapitulated in the in situ Hi-C and intact Hi-C data, with the latter two also exhibiting stripes and TADs (**Figure 4a**, bottom). To answer whether such conservation of jets across technologies is common, we computed the overlap between Repli-HiC and in situ or intact Hi-C. Many of in situ Hi-C (122 of 312) and intact Hi-C (113 of 274) jets were also identified in Repli-HiC data, corresponding to 39% and 41% for in situ Hi-C and intact Hi-C respectively (**Figure S4a**).

**Figure 4:**
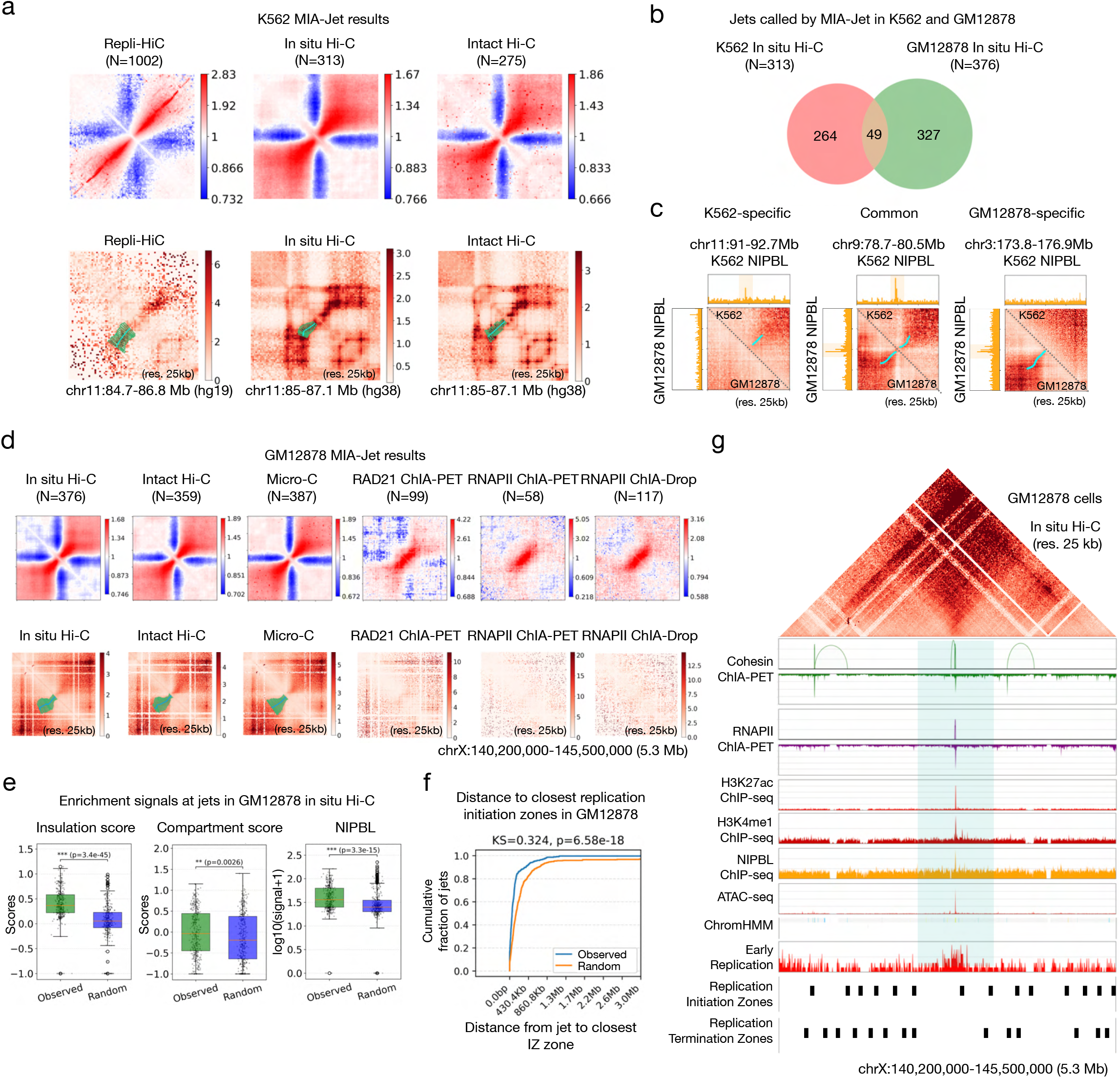
MIA-Jet results in K562 and GM12878 human cell lines. **(a)** Top: Aggregate Observed/Expected maps of jets identified by MIA-Jet in K562 Repli-HiC, in situ Hi-C, and intact Hi-C. Bottom: Example location of a jet identified in all three datasets with MIA-Jet annotations in lower triangle. **(b)** Venn diagram comparing jets in K562 in situ Hi-C and GM12878 in situ Hi-C. **(c)** Left: Example jet location called in K562 but not GM12878; K562 contact map and NIPBL track on upper right and GM12878 contact map and NIPBL track on lower left. Middle: Example jet location called in both K562 and GM12878. Right: Example jet location called in GM12878 but not K562. Contact maps are Observed/Expected. **(d)** Top: Aggregate Observed/Expected maps of jets identified by MIA-Jet in GM12878 cells for 6 datasets: in situ Hi-C, intact Hi-C, Micro-C, RAD21 ChIA-PET, RNAPII (RNA Polymerase II) ChIA-PET, and RNAPII ChIA-Drop. Bottom: 5.3 Mb region on chromosome X with MIA-Jet annotations in lower triangle, if detected. **(e)** Boxplots of insulation score, compartment score, and log-transformed NIPBL ChIP-seq. Observed: 376 jets identified by MIA-Jet in GM12878 in situ Hi-C data. Random: 376 random regions (**Methods**, *Generating random control*). **(f)** Empirical cumulative distribution function of distance from “Observed” or “Random” locations to closest replication initiation zone. Observed: 376 jets identified by MIA-Jet in GM12878 in situ Hi-C data. Random: 376 random regions (**Methods**, *Generating random control*). P-values from two-sided Kolmogorov-Smirnov test. **(g)** The same 5.3 Mb jet region as presented in bottom of panel **(d)**, with GM12878 in situ Hi-C Observed/Expected contacts on top, loops and binding peaks of cohesin and RNAPII ChIA-PET below. The binding profiles for H3K27ac, H3K4me1, NIPBL ChIP-seq, and ATAC-seq are displayed below, with chromHMM state denoting enhancer (yellow) at the jet origin. Bottom three tracks show early repli-seq data, replication initiation zones, and replication termination zones. The jet region identified by MIA-Jet is highlighted in light blue.

By contrast, the jets identified in GM12878 cells generally did not overlap those in K562 cells using the same in situ Hi-C technique, with only 49 common jets between them and a Jaccard index of 0.08 (**Figure 4b**, top panel). However, despite their small number, the common jets showed signal strengths comparable to the K562-specific and GM12878-specific jets (**Figure S4b**), implying a small, yet prominent set of jets shared between the two cell lines. This prompted the natural question of whether cohesin loading (NIPBL) at these genomic locations correlates with the presence or absence of jets. Exemplary loci suggest that a K562-specific jet is supported by NIPBL signal that is lacking in GM12878, while GM12878-specific jet had strong NIPBL binding that is lacking in K562 (**Figure 4c**, left, right). In keeping with this line of reasoning, we expect a common jet between K562 and GM12878 to have high NIPBL signal in both their respective NIPBL tracks, as shown in the example (**Figure 4c**, middle). Genome-wide statistics indicate that the number of GM12878 NIPBL peaks are indeed much higher at the jet origins of common (2.49) and GM12878-specific (2.27) jets than at K562-specific sites (1.27) (**Figure S4c**, left). Similarly, K562 NIPBL peaks were on average higher in K562-speicific (1.91) and common (2.94) sites than at GM12878-specific jet origins (**Figure S4c**, right). These results suggests that while jets are not widely shared among GM12878 and K562, among the jets that are common, there is strong jet signal at shared NIPBL binding sites. Beyond NIPBL, we next sought to identify other genomic features associated with jets, leveraging the publicly available datasets and diverse 3D genome mapping technologies profiled for the GM12878 cell line.

### Jets are enriched at cohesin loading sites and are associated with early replication

We first asked whether enriching for chromatin interactions mediated by protein factors retains jets present in Hi-C experiments. In particular, we chose a cohesin subunit RAD21 as jets may arise from cohesin extrusion, and RNA Polymerase II (RNAPII) as jets were shown to be enriched at active enhancers. While RAD21 ChIA-PET, RNAPII ChIA-PET, RNAPII ChIA-Drop data exhibit jet-like signals (**Figure 4d**, top panel), their jets are fewer in number (58-117) compared to Hi-C (376). At a strong jet location on chromosome X, MIA-Jet identified diffused jets in Micro-C, in situ Hi-C, and intact Hi-C, but did not report any jets in RAD21 ChIA-PET, RNAPII ChIA-PET and RNAPII ChIA-Drop data, implying that RNAPII may not be involved in forming this particular jet (**Figure 4d**, bottom panel). Computing overlap of jets between all pairs of datasets shows in situ Hi-C, Micro-C and intact Hi-C to have the highest number of overlapped jets, followed by overlaps among the ChIA-PET and ChIA-Drop datasets (**Figure S4d**).

We next used jets identified by MIA-Jet (‘Observed’) to characterize their genomic features compared to random regions (‘Random’) (**Methods**, *Generating Random Control*). Across many platforms, jets had higher insulation and compartment scores than random regions (**Figure 4e, S4e, S4f**), potentially suggesting that jets reside in active regions within TADs. As shown in mouse DP thymocytes, human GM12878 cells also had jets with enriched NIPBL binding signals (**Figure 4e, S4g**) which may coincide with cohesin loading sites related to extrusion activities.

Based on these observations, we focused on the 376 jets identified in the GM12878 in situ Hi-C data as it does not have any bias towards protein-enriched or replication-enriched activities (e.g., ChIA-PET or Repli-HiC). While these epigenomic features hint that jet origins are likely cohesin loading sites, as was assumed by Guo et al. in concluding that jets arise due to cohesin extrusion, we adopted a more rigorous definition of true loading sites. In addition to having high 1-dimensional binding of active marks and NIPBL, we impose that cohesin loading sites are not bound by CTCF, far from nearest CTCF binding sites, and is encapsulated by at least one strong convergent CTCF loop (see **Methods**, *Defining GM12878 cohesin loading sites and anchor sites*). Doing so revealed that jets are more likely to originate from cohesin loading than random regions but explains only 58 of 376 jets (**Figure S4h**). Chromatin states at jet origins were mostly enhancers, promoters or transcribed states (303 of 376 jets), while random regions only had 214 enhancer/promoter marks and were largely heterochromatin (**Figure S4i**). Interestingly, Hi-C jets were significantly closer to replication initiation zones (IZ) than the random control (**Figure 4f**), which is consistent with Type I jets observed in Repli-HiC data on K562 cell line (Liu et al., 2024). When we examined the jet shown in **Figure 4g**, the jet origin shows binding of cohesin (with small loops) and RNAPII in ChIA-PET data. This region is also enriched in H3K27ac, H3K4me1, NIPBL, and ATAC-seq signals as well as being annotated as active enhancer by chromHMM (**Figure 4g**). Repli-seq data indicates that the jet origin corresponds to early DNA replication signals. In addition, a replication initiation zone aligns with the jet origin, which is not the case for replication termination zones that instead pepper the flanking regions near the jet extrusion point.

### Depletion studies in human colorectal cancer cells reveal jets are broadly eliminated without cohesin

The auxin-inducible degron (AID) system (Natsume et al., 2016) is a powerful tool for rapidly depleting protein factors for studying their function *in vivo*. Applying these tools to human HCT116 colorectal cancer cells revealed that depleting RAD21 cohesin sub-unit results in loss of all chromatin loops (Rao et al., 2017). However, little is known about the effect of cohesin degradation on chromatin jets in this cell line, mainly due to the lack of computational tools to identify and quantify jets genome-wide. Here, we applied MIA-Jet to identify jets in the same dataset.

In a region on chromosome 13, a jet was identified in Hi-C data before depleting RAD21 (‘0hr’) but the jet structure disappears in the Hi-C map after 6 hours of auxin treatment for RAD21 depletion (‘6hr’) (**Figure 5a**). As was the case with GM12878 jets (**Figure 4e**), the jet origin is enriched for positive insulation score, but only before cohesin depletion. As for replication signals, publicly available 16 fraction repli-seq data on HCT116 cells (Zhao et al., 2020) shows the surrounding jet region with higher replication signal at early stages in S phase (P03-P08) compared to later (P09-P17), consistent with the early replication IZ annotation and the dual-stage (early, late) repli-seq data (**Figure 5a**).

**Figure 5:**
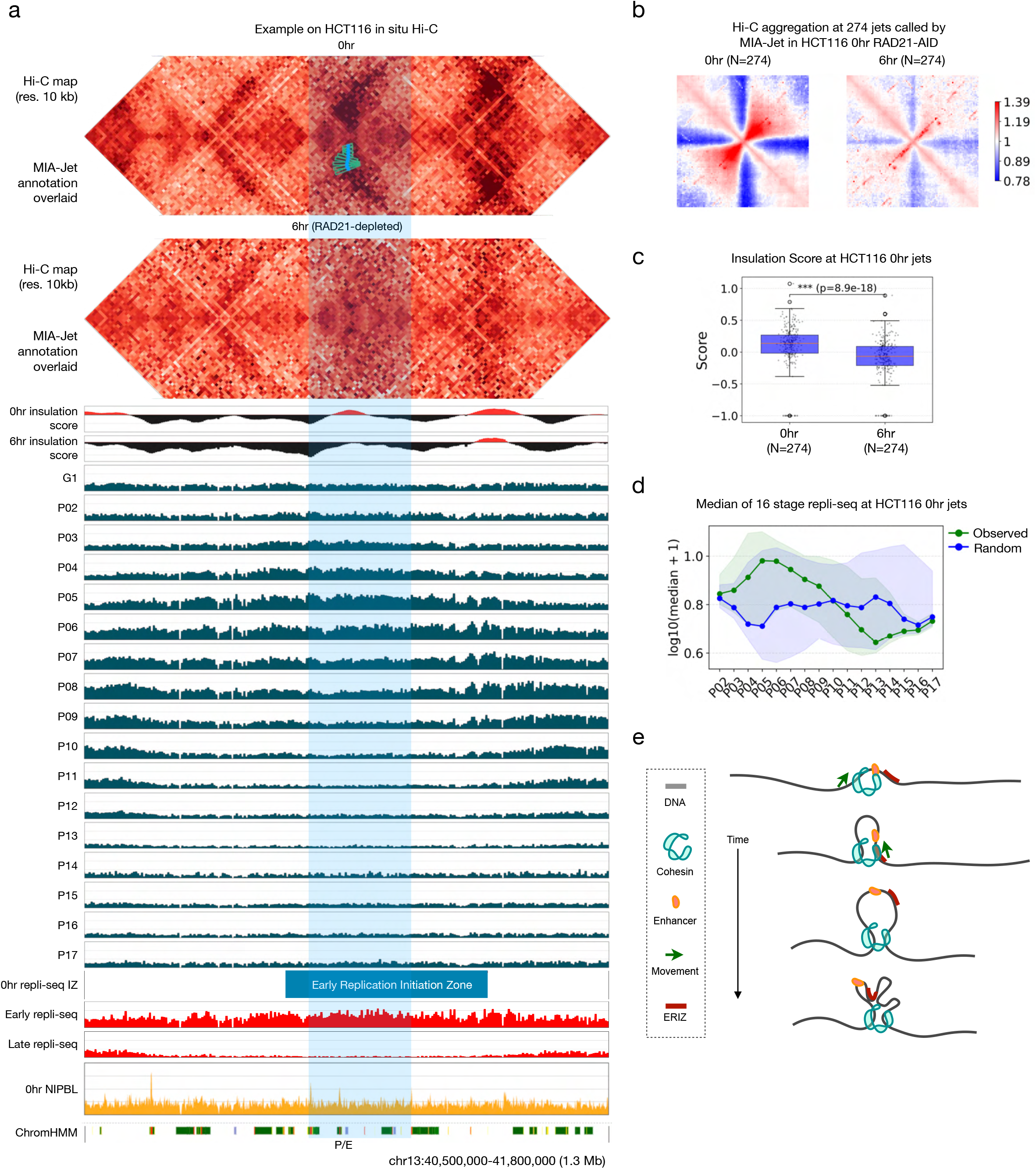
Effects of cohesin depletion on chromatin jets. **(a)** Data shown for HCT116 RAD21-mAC cell line engineered to rapidly deplete RAD21 with auxin treatment. ‘0h’: without auxin treatment (before depletion; control), ‘6h’: 6 hours of auxin treatment (RAD21 depletion). A region encompassing the extrusion point of a jet called by MIA-Jet is highlighted in light blue. Top two tracks show 0hr and 6hr insulation score. G1 repli-seq data and 16 S-phase fractions P02-P17 is shown with same y axis limit, followed by 0hr repli-seq initiation zones, early and late repli-seq tracks, 0hr NIPBL ChIP-seq and ChromHMM states denoting enhancers (yellow), promoters (red) and transcription (green). **(b)** Aggregate Observed/Expected map of jets identified by MIA-Jet in HCT116 0hr (left) and aggregating the same locations in HCT116 6hr data (right). **(c)** Boxplot of 0hr and 6hr insulation score at jets identified by MIA-Jet in HCT116 0hr. **(d)** Median of HCT116 16-fraction repli-seq data. Observed: 274 jets identified by MIA-Jet in HCT116 0hr. Random: 274 random regions (**Methods**, *Generating random control*). Shaded contour indicates 25^th^ and 75^th^ percentiles. **(e)** A proposed model of chromatin jet formation, where cohesin first loads onto the DNA at active enhancer sites near early replication initiation zone (ERIZ), then alternately extrudes from the left and right strands until the extruded strands fold among themselves.

For genome-wide analysis, we ran MIA-Jet on 0hr and 6hr RAD21-depleted HCT116 Hi-C data, where we observed a two-fold reduction in the number of jets: 274 at 0hr and 137 at 6hr (**Figure S5a**). Aggregate map after cohesin depletion showed spurious high value signals mixed with a weaker jet profile as compared to the untreated cells (**Figure S5a**). Overlaps indicated that only a small number 27 out of 274 jets persisted after cohesin depletion (**Figure S5b**). To observe the effects of cohesin depletion on the jet strength, we aggregated the 274 jet locations identified before cohesin depletion and show the 6hr depleted Hi-C maps at the corresponding 274 locations, which revealed significant reduction in the strength of jets (**Figure 5b**). Focusing on these 274 jet regions before cohesin depletion, we saw positive insulation score across the genome, while the scores were markedly lower in RAD21-depleted cells (**Figure 5c**). In line with the example HCT116 jet which showed higher replication signal at early S phase stages (**Figure 5a**), genome-wide jet locations show a similar trend: a gradual increase followed by a decrease in replication signals across S-phase, with a peak occurring at stage P05 and P06 (**Figure 5d**). Inspecting jet origins for replication initiation zones (IZs) revealed a two-fold increase in the number of ‘early’ replication IZ (ERIZ) (80) compared to the random control (39) (**Figure S5c**). We then asked whether this enrichment with early replication signal is specific for the 0hr jets or also present with the 6hr jets by computing the genomic distance from the jet origin to the closest ERIZ. We found that 0hr jets were significantly closer (p-value 1.25e-5) to ERIZs compared to the random control, whereas 6hr jets were not (p-value 0.629) (**Figure S5d**).

These results confirm that the high insulation score, NIPBL binding, enhancer or promoter chromatin states, and early replication initiation zone in GM12878 in situ Hi-C jets are recapitulated in the HCT116 in situ Hi-C jets. Furthermore, the prevalent depletion of jets in RAD21 depletion HCT116 cells suggests the potential role of cohesin in forming jets.

A motivation for developing MIA-Jet was to determine whether jets arise as part of cohesin-CTCF loop extrusion model or as DNA replication. However, jets in HCT116 and GM12878 cell lines supports both scenarios: jets are enriched at active, cohesin loading sites, and proximal to replication initiation zones, with a high insulation score revealing that these regions are brought close together in 3D space. Jet origins are thus highly dense clusters of transcriptional and replication-associated marks that occupies a relatively small interval on the chromatin strand. One interpretation is that cohesin extrusion and DNA replication processes are interdependent on each other, but such hypothesis requires additional experimental validations. We posit that cohesin loads onto the DNA at localized active sites, and extrudes in a slow, two-sided asymmetric manner, sometimes reeling the right strand or the left strand (**Figure 5e**). It is unclear whether the enriched chromatin marks near jets (enhancer states, early replication marks, or even CTCF boundary elements) influence the slowing of the jet extrusion process. Moreover, the potential role of other protein factors in forming or maintaining jets remains elusive, as are the generalizability of their roles across various species. To answer these questions, we focused on publicly available 3C data for additional protein depletions.

### Cohesin is required for jet formation, whereas CTCF shapes jet extrusion trajectory

We first tested whether MIA-Jet can call jets in non-mammalian cell lines, namely in zebrafish embryos where “fountains” were previously reported (Galitsyna et al., 2026). MIA-Jet could annotate an example jet region and capture its broadening width (**Figure S6a**, left), which is consistent with the genome-wide aggregate maps (**Figure S6a**, right). We then applied MIA-Jet to *C. elegans* in situ Hi-C data where recent studies have found fountains that have been largely eliminated upon a depletion of SMC-3, a subunit of cohesin, and extended with WAPL-1 depletion (Kim et al., 2025). As expected, MIA-Jet results conform to this pattern. Aggregate maps show reduced jet strength with SMC3 depletion and stronger jets for WAPL depletion compared to the control (**Figure 6a**). Aggregating the 109 jet locations called in control with the depletion Hi-C data confirms that jets at the same location are attenuated upon SMC3 depletion and their signals are stronger without WAPL (**Figure S6b**). Of these 109 jets called in control, 76% were identified in the WAPL-depleted data, suggesting that MIA-Jet can identify many WAPL-dependent jets from the control cell line (**Figure S6c**). Example jet on chromosome IV highlights a jet that was called in control and WAPL-depleted maps but not in the SMC3-depleted data, with the jet annotation reflecting a longer and slightly curved jet shape after WAPL depletion (**Figure 6b**).

**Figure 6:**
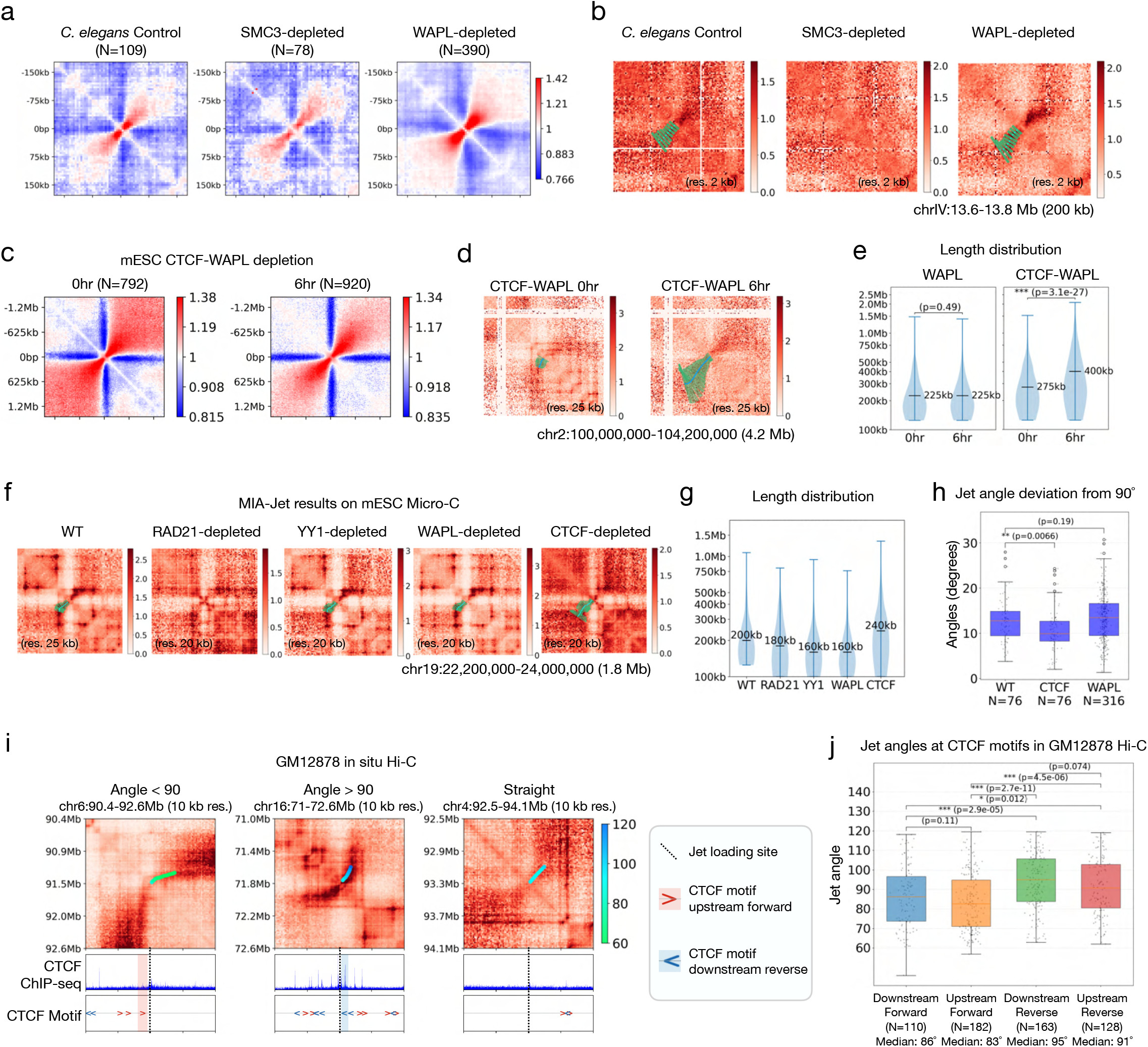
The role of CTCF and WAPL in jet formation. **(a)** Aggregate Observed/Expected maps of jets identified by MIA-Jet in *C. elegans* in situ Hi-C data: control (left), SMC-3 depletion (middle), and WAPL-depletion (right). **(b)** Example jet in *C. elegans* with MIA-Jet annotation, if called. **(c)** Aggregate Observed/Expected maps of jets identified by MIA-Jet in mESC in situ Hi-C data: 0hr and 6hr CTCF-WAPL depletion. **(d)** Example jet in 0hr and 6hr mESC CTCF-WAPL depletion with MIA-Jet annotation. **(e)** Length distributions of 0hr, 6hr WAPL depletion and 0hr, 6hr CTCF-WAPL depletion. Median length and p-values for two-sided Wilcoxon rank-sum test are shown. **(f)** An example of curved jet in mESC Micro-C data (WT) and protein (RAD21, YY1, WAPL, CTCF) depleted cells, with MIA-Jet annotation in lower triangle. **(g)** Length distributions showing median length of jets called in WT, RAD21, YY1, WAPL, and CTCF protein depleted micro-C data. **(h)** Mean deviation of jet angles from 90° using MIA-Jet annotations. WT, CTCF: Only the common 76 jet locations between WT and CTCF-depleted maps are shown. WAPL: All 316 jets called in WAPL-depleted is shown. **(i)** Example jets with MIA-Jet line annotation coloured by angles in GM12878 in situ Hi-C data. Left: jet angled less than 90° with corresponding CTCF motif upstream and forward orientation in red highlight. Middle: jet angled greater than 90° with corresponding CTCF motif downstream and reverse orientation in blue highlight. Right: straight jet with no intersecting CTCF motifs. **(j)** Boxplots of mean jet angle corresponding to CTCF motifs (downstream forward, upstream forward, downstream reverse, and upstream reverse) for GM12878 Hi-C data. N is the number of jets intersecting with each CTCF motif category. P-values from two-sided Wilcoxon rank-sum test.

An important difference between the mammalian genome and *C. elegans* is that the latter lacks CTCF or functional homolog (Kim et al., 2025; Luthi et al., 2025). Thus, while it is clear that CTCF is not necessary for the formation of jets in *C. elegans*, we asked the same question – whether CTCF is necessary in mammalian cell lines, and if not, what influence, if any, it has on jet formation. To address this question, we applied MIA-Jet on mouse embryonic stem cells (mESC) in situ Hi-C data with 6hr WAPL depletion. Across the genome, MIA-Jet results show a weaker enrichment in the upper right corner of the aggregate plots for the 6hr depleted WAPL data compared to 0hr control (**Figure S6d**, left). Inspecting an example region chr2:100-104.2Mb, we find that WAPL depletion increased the strengths of stripes or loops (**Figure S6d**, right) in line with the results from a previous WAPL-depleted study in human HAP1 cells (Haarhuis et al., 2017). In stark contrast, MIA-Jet in CTCF-WAPL double depletion data reveals an aggregate map with an elongated jet shape (**Figure 6c**), more so than the WAPL only depletion (**Figure S6d, S6e**). Indeed, the same example region chr2:100-104.2Mb as shown before has a lack of stripes and loops upon CTCF-WAPL double depletion, and a large jet (4Mb) in the 6hr CTCF-WAPL depleted map (**Figure 6d**). The likely explanation is that depleting WAPL cannot create extended jets while the CTCF boundary element is present. This is supported by genome-wide length distributions, where we find a significant increase in jet lengths with 6hr CTCF-WAPL depleted (median = 400kb) compared to 0hr (median = 275kb) (**Figure 6e**, right), but not when only WAPL is depleted (**Figure 6e**, left). Indeed, loops or stripes are formed largely at CTCF binding sites with the current viewpoint that CTCF blocks cohesin from extruding (Rao et al., 2014; Vian et al., 2018). With increased cohesin rings residing on the DNA due to the lack of its unloading factor WAPL, there is expected to be only more stalling at these sites (i.e., increased enrichment of stripes and loops) compared to control (**Figure S6d**). Removing both CTCF and WAPL in mESCs recreates the conditions in WAPL-only depletion in *C. elegans*, wherein neither CTCF nor WAPL is present, resulting in extended jets.

Finally, we applied MIA-Jet to Micro-C mESC data with extended protein depletions: RAD21, YY1, WAPL, CTCF, and control. In the control, a clear curved jet is identified by MIA-Jet (**Figure 6f**). However, this structure is attenuated upon RAD21 depletion, resulting in no jet found by MIA-Jet. By contrast, depleting YY1 had minimal effect on the jet, while WAPL depletion resulted in a map with additional loops but minimal change to the jet length (**Figure 6f**). CTCF-depleted map shows weaker loops, in which MIA-Jet identified a jet with increased length and with a straight, symmetric direction of extrusion than the curved jet seen in WT (**Figure 6f**). We aggregated the 240 jets observed in mESC WT cell line and confirmed large depletion in signal with RAD21 whereas YY1 and WAPL were generally unaffected (**Figure S6f**). Genome-wide annotations support the example: depleting CTCF resulted in longer jets (median = 240 kb) compared to WT (median = 200 kb) (**Figure 6g**). Furthermore, we used the jet angles from MIA-Jet to compute how far each jet deviates from a straight (90°) jet, which allowed us to observe that depleting CTCF resulted in significantly less angled jets than the WT (**Figure 6h**). In line with the results from WAPL depletion data in *C. elegans*, we reasoned that although CTCF may not be necessary for jet formation, it may stop or slow jet formation in a manner that is similar to stripes (i.e., asymmetric CTCF blocking on one side of cohesin extrusion). For instance, exemplary regions in GM12878 in situ Hi-C data show a CTCF motif upstream (left of dotted line) of jet origin in a forward (>) orientation is correlated with a jet curved to the right with angle < 90 degrees (**Figure 6i**, left panel). Similarly, a jet curved to the left with angle > 90 degrees is accompanied by a CTCF motif downstream (right of dotted line) in a reverse (<) orientation (**Figure 6i**, middle panel). Such bias is not observed in a case where there is no CTCF binding nearby straight jet (**Figure 6i**, right panel).

We sought out to rigorously test this reasoning: whether a jet that encounters a CTCF protein slows the speed of extrusion on one side, causing the jet to curve in a certain orientation and whether this process is dependent on the relative location with the jet origin and CTCF binding motif orientation. Specifically, for each CTCF motif that encounters a jet, we tested whether the corresponding jet angle is affected, where CTCF motifs may encounter a jet either downstream or upstream of the jet loading site. If the CTCF motif is downstream (**Figure S6g**, left), we look at the x-axis jet coordinates and their corresponding angles from MIA-Jet. If the CTCF motif is upstream (**Figure S6g**, right), we use the y-axis jet coordinates and their corresponding angles from MIA-Jet. We performed genome-wide statistics of all four possible cases of CTCF location (CTCF motif that is *downstream* or *upstream* of the jet loading site) and CTCF motif orientation (*forward* or *reverse*). For each case, we extract the corresponding jet angle according to the schematic (**Figure S6g**), which we plot as a boxplot for each category (**Figure 6j**). As seen in the example (**Figure 6i**), genome-wide statistics indicate that the jet angles for the “Upstream Forward” category is less than 90° (median = 83°) and that for “Downstream Reverse”, the angle is greater than 90° (median = 95°) with a significant difference between them (p-value = 2.7e-11) (**Figure 6j**). This pattern is not observed with “Upstream Reverse” (median = 91°) and “Downstream Forward” (median = 86°) with weak significance (p-value = 0.012), suggesting that the orientation of the CTCF motif is critical in affecting the direction of jet extrusion (**Figure 6j**). We repeated this analysis for K562 in situ Hi-C data and observed similar patterns (**Figure S6h, S6i**). Altogether, we find that CTCF motifs that are oriented towards the jet origin—often cohesin loading sites—has a greater propensity to affect jet extrusion than motifs that are oriented away from the loading site.

## DISCUSSION

We described MIA-Jet and demonstrated its utility in accurately identifying jets of various diffuseness, lengths, widths, and angles. Notably, it does not require training steps or kernels, thereby allowing users to detect jets in a wide range of settings in 3D genome mapping data. MIA-Jet is also one of the few publicly available software for calling jets. Recently, an improved version of the Fun algorithm, called Fun2, and a high-resolution replication-associated chromatin interactions data Repli-MiC identified angled jets (Zhangding et al., 2026), but Fun2 was not designed to identify jets with variable curvatures as MIA-Jet identified in various Hi-C datasets with and without protein depletion (**Figures 1b, 6f, 6i**). Similarly, Rep-Hi-C experiments revealed that the fountain strength increased under replication stress (Gaggioli et al., 2026).

These experimental advancements to probe chromatin interactions associated with DNA replication highlight the critical need for a versatile computational algorithm that can not only identify but also quantify various features of jets. MIA-Jet represents a first step towards meeting this need, yet there are limitations of MIA-Jet in its current form. Of note, MIA-Jet is dependent on several parameters that have been finetuned for each genomic technologies and organisms, such as the resolution and window size. While MIA-Jet outperformed Fun and Fontanka in simulated data with robust F1 scores across various resolution, window size, jet width, and q-value (**Figures 2b, 2c**), it was sensitive to sequencing depth especially at 10% of the original reads (**Figure S3d**).

The MIA-Jet findings offer answers to several biological questions yet also pose new ones. Our motivation for developing MIA-Jet was to determine whether jets arise as part of the two-sided cohesin loop extrusion model or as DNA replication. However, to our surprise (and slight disappointment), the results supported both scenarios: jets were enriched at cohesin loading sites (**Figures 3d, 4g**) as well as early replication initiation zones (ERIZ) (**Figures 4f, 4g**). One interpretation is that cohesin extrusion and DNA replication processes are interdependent on each other and thus jets may arise due both cohesin-mediated loop extrusion and replisome-led replication activities. In particular, Repli-MiC data showed that fountains at the transcription start site of active genes are angled towards the direction of transcription (Zhangding et al., 2026), akin to cohesin ChIA-Drop contacts being biased towards the transcription orientation alongside RNA Polymerase II (Kim et al., 2026) and cohesin ‘islands’ suggesting that transcription can translocate cohesin. These reports hint at the intricate interplay among cohesin, RNA Polymerase II, and replisome to perform replication and transcription in 3-dimensional space, but such hypothesis requires additional experimental validations. We posit that cohesin loads onto the DNA near the ERIZ and extrudes in a two-sided asymmetric manner—sometimes reeling the right strand or the left strand—and the extruded DNA strands crumble up to form a TAD-like structure (**Figure 5e**). Intriguingly, it is unclear which environmental forces or protein factors stops jet extrusion, analogous to the role of convergent CTCF on stopping chromatin loops. In addition, jets seem to gradually fade and increase in diffuseness as they move away from the main diagonal. Both the underlying molecular mechanism and biophysical principles for such patterns are yet to be explored. By leveraging publicly available Hi-C data, one can also ask if certain cell types encompass more jets, and whether jets are conserved across cell lineages or are cell-type-specific. Integrative analyses of all chromatin structures, ranging from large-scale TADs and compartments to finer-scale loops, stripes, and jets would be worthwhile. As the 3D genomics field ventures into this new area of chromatin biology, we envision MIA-Jet to provide the important first steps towards understanding biological implications of chromatin jets and beyond.

## METHODS

### Notation

Let ℝ denote the reals. Unless otherwise stated, we reserve capital bold face (e.g. **X** ∈ ℝ^*M*×*N*^) for matrices, script (e.g. *Y*) for sets, capital letters e.g. (*M*) to refer to matrix dimensions, and lower case for scalars (e.g. *x*). Let the *i*jth coordinate of **X** be denoted as **X**(*i*, j). Let X(*i*, : ) refer to the *i*th row of **X** and **X** (:, j) refer to the jth column of **X**. Let the eigenvalues of an *N* × *N* matrix **X** ∈ ℝ^*N*×*N*^ be *λ*_1_(**X**), *λ*_2_(**X**), …, *λ*_*N*_(**X**) where *λ*_1_(**X**) ≥ *λ*_2_(**X**) … ≥ *λ*_*N*_(**X**). Let *L⋅*], ]*⋅*] be the floor and ceiling function, respectively.

### Inputs to the program

At a minimum, the MIA-Jet program (v2.0.3) takes as input the path to the .hic file, the chromosome, the Hi-C resolution, and the experiment type `exp_type`, which is either “hic” (default) or “replihic”. The experiment type is designed to automatically assign default parameter values depending on whether the data is from a Hi-C experiment or the Repli-HiC experiment (Liu et al., 2024), summarized in **Supplementary Table S1**, ‘MIA-Jet Parameters’. The reasoning behind this is that jets differ in their shape and length depending on the experiment; Hi-C jets reported in mouse DP thymocytes (Guo et al., 2022) are thicker and have greater variation in jet thickness and angle than Repli-HiC jets, which are comparatively thinner and exhibit significant variation in length (e.g., Type I and Type II jets). In particular, MIA-Jet can identify jets that curve in particular directions. The user parameter `angle_range` specifies the lower bound and upper bound for the expected jet angles. A straight jet that is perpendicular to the main diagonal is 90°, a vertical stripe is 45° and horizontal stripe is 135°. Setting the `angle_range` to 45-135 will capture some vertical and horizontal stripes; a smaller range, such as 80-100 will capture straight jets. The default range is 60-120 if `exp_type`=‘hic’, 80-100 if `exp_type`=‘replihic’. In addition to differences with the angle, Hi-C contact maps contain many other structures such as stripes, loops, and A/B compartments, requiring additional splitting and filtering criteria for Hi-C.

Default parameter values have been set to optimize recovering jets seen in WT DP Thymocytes at 25 kb resolution (Guo et al., 2022) for `hic` mode and Repli-HiC jets at 25 kb resolution (Liu et al., 2024) in `replihic` mode. Full parameter specifications (v2.0.3) are detailed in the **Supplementary Table S1** ‘Analysis Parameters’.

### Generating an Image from a Hi-C file

#### Processing Hi-C Data

Let the contact matrix of chromatin interaction frequencies be **A** ∈ ℝ^*P*×*P*^, where the entry A(*i*, j) is the interaction frequency between genomic loci *i* and j. The contact matrix **A** is read with the *hic-straw* package (1.3.1) using the user-specified `data_type`, `normalization`, and `resolution`. Rows and columns that sum to 0 are identified and removed. Let ε *⊆* {1,2, …, *P*} be this set of excluded indices and define 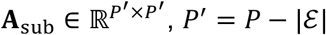, as the matrix with rows and columns that sum to 0 removed. We remove these regions as kernel convolution produces artefacts if there is a large discrepancy in image intensity values. For the `data_type` parameter (default ‘oe’ if `exp_type`=‘hic’, ‘observed’ if `exp_type`=‘replihic’), MIA-Jet can work with both “observed” (untransformed Hi-C values) and “oe” (Observed over Expected, which divides each off-diagonal by its mean). If the `data_type` is “observed” we apply a log-transform to reduce the dynamic range from the distance-dependent decay. The contact matrix is min–max normalized to [0,1] with `normalize` from *opencv* (4.11.0), which we denote as **A**_I_.

#### Generating the Hi-C Image

We extract a rectangular box in the upper triangle of **A**_I_, rotated such that the base of the rectangle aligns with the main diagonal of **A**_I_ to yield a rectangular image **I** ∈ ℝ^*M*×*N*^, with width (number of columns) and height (number of rows) given by

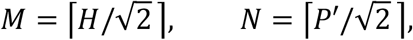

where *H* is the `window_size` user parameter in bins. This is done by rotating the image 45° using `rotate` from *scipy* (1.16.1) with boundary mode set by the `rotation_padding` parameter (default ‘nearest’). From hereon we refer to the rotated rectangular image I as the *Hi-C Image* (see **Figure S1** ‘Hi-C image generation’). We extract a rectangle to improve computational efficiency as A_+_ is a *P*^′^ × *P*^′^ matrix, where *P*^′^ is large at high resolutions (e.g., 5 kb). Running convolution operations on the entire matrix is unnecessary as the matrix is symmetric, so we extract a rectangle box in the upper triangle. Also, since jets are typically found near the main diagonal (see Liu et al., 2024, Guo et al., 2022), we do not need the entire upper triangle of **A**, motivating the binned window size parameter *H*.

### Computing scale space features

#### Basic Principles of Scale Space Ridge Detection

Scale space has been used in image processing to identify structures of interest appearing at various scales (Witkin, 1984). The basic idea is to convolve an image with a Gaussian kernel at a continuum of standard deviations, generating a stack of images that are progressively smoothed versions of the original image. At one extreme, if the standard deviation is 0 then no blurring is applied to the image such that fine-scale objects are present. As we increase the blur amount, the image loses fine-scale detail and keeps only larger scale objects. Thus, the amount of blur affects the scale of structures we wish to capture. The family of blurred images is the “scale space” of the image and can be used to identify the appropriate scale of particular objects or features.

We first motivate scale space by describing the object of interest, which are jets. A jet is an elongated streak of elevated contact frequency emanating from the main diagonal (**Figure 1a**). If we view the Hi-C image as a surface, where the coordinate plane is indexed by (*i, j*) and the surface by the contact intensity **I**(*i*, j), then one can imagine it is possible to walk along the jet, where the surface to the left and right is lower than the surface we are on. Mathematically, this corresponds to a local maximum perpendicular to the direction of the jet, or a negative second directional derivative. Indeed, the more locally enriched the jet appears (stronger jet), the more negative the second directional derivative. This quantity is the largest magnitude eigenvalue of the Hessian, which we use an indicator of jet strength (ridge strength) **Λ**. The corresponding eigenvector gives the direction **Θ**(ridge angle) that the jet is heading towards (see *Feature Generation*; **Figure 1d** ‘Feature generation’). Both the definition and mathematical characterization of this recently discovered chromatin jet is not new and has been well studied as a *ridge* in image processing (Haralick, Watson and Laffey, 1983).

Importantly, a jet has a width associated to each point of the jet, defined as the extent to which the left and right sections slope away. For instance, a jet may be narrow near its root at the diagonal and broaden as it extends away. This is harder to compute and other works have provided methods to infer the widths (Steger, 1998). Here, we use automatic scale selection (Lindeberg, 1998) to assign widths to jets. At a high level overview, we convolve the Hi-C image with a Gaussian kernel of increasing standard deviation *s*, producing a family of smoothed images **I**^(*s*)^ parametrized by *s*; for each blurred image we then compute the ridge strength map using normalized derivative kernels obtaining **Λ**^(*s*)^. Then, the appropriate scale for each point of a jet can be obtained by finding the local maxima of **Λ**^(*s*)^ with respect to scale *s* (see *Feature Generation* and **Figure S1** ‘Scale inference’). A similar scale-space method has been used in Hi-C data analysis that uses the Laplacian to detect blob-like structures (loops) at various scales (Roayaei Ardakany et al., 2020).

#### Gaussian Blurring

Let T = (*s*_1_, *s*_2_, …, *s*_*d*_) denote the tuple of standard deviations of the Gaussian kernels used in scale space, ordered such that *s*_1_ ≤ *s*_2_, ≤ *⋯* ≤ *s*_*d*_. Let the index function return the index of the scale (e.g., index(*s*_3_) = 3). Let I^(*s*)^ ∈ ℝ^*M*×*N*^ denote the image blurred by a 2D Gaussian kernel X_*s*_ with scale *s ∈* {*s*_1_, *s*_2_, …, *s*_*d*_} i.e.,

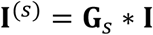

where *∗* is the convolution operator. For a blurred image **I**^(*s*)^, the basic principle of scale space is that if *s* is small then we can find narrow jets. Conversely, a larger *s* enables the detection of wider jets. From hereon we refer to each standard deviation *s ∈* {*s*_1_, *s*_2_, …, *s*_*d*_} as a *scale*. The jet width is the cross-sectional distance perpendicular to the extrusion direction (in number of pixels). Each point of a jet may have a different width. For instance, a jet that gradually becomes broader as it extrudes has a smaller width near the root than at points further away from the main diagonal. If we want to capture such jets, the set of scales {*s*_1_, *s*_2_, …, *s*_*d*_} must be chosen such that the minimum scale and maximum scale cover the range of possible jet widths to detect. Alternatively, users may also specify this range directly via the `jet_widths` parameter, which is the lower and upper bound in pixels of the desired jet widths.

#### Choosing the scales

Let the `jet_widths` be denoted as w_*lb*_, w_*ub*_, (w_*lb*_ < w_*ub*_) which denotes the smallest (lower bound, ‘lb’) and largest (upper bound, ‘ub’) jet widths (in pixels) the program should detect (default `None`). The width range w_*lb*_, w_*ub*_ is converted to scales according to the following formula from ImageJ Ridge Detection (https://imagej.net/plugins/ridge-detection):

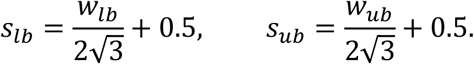

We then sample between *s*_*lb*_ and *s*_*ub*_ in log scale, following the scale space framework (Koenderink, 1984):

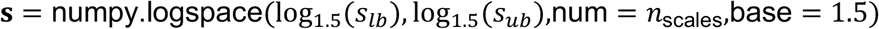

where the number of scales

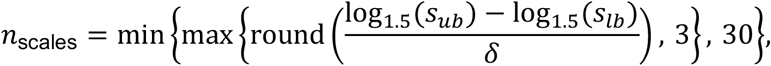

provides a reasonable number of scales (minimum 3 and maximum 30) in log-space with δ = 0.25 defining the sampling resolution. Since automatic scale selection involves computing local maxima with respect to scale (*Scale inference*), we impose a minimum of 3 scales for the finite difference. The maximum on the number of scales is imposed for computational efficiency.

Alternatively, the `scale_range` parameter (default `None`) enables users to specify set of scales {*s*_1_, *s*_2_, …, *s*_*d*_} directly. If neither `jet_widths` or `scale_range` is given, the default grid is used,

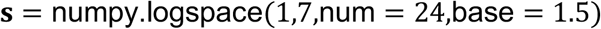

which corresponds to detecting jet widths that can be 3 to 60 pixels wide – a range that may be extreme for most applications. If possible, it is recommended to visualize the Hi-C maps and count roughly how wide the jets are in pixels and use these to specify the lower and upper bounds via the `jet_widths` parameter.

Finally, there may be scales *s ∈* {*s*_1_, *s*_2_, …, *s*_*d*_} that are too “large” for the image dimensions, which arises when the Gaussian kernel size is too large to convolve with the image. Any scale *s ∈* {*s*_1_, *s*_2_, …, *s*_*d*_} such that 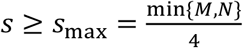 are resultingly removed.

#### Feature Generation

For each scale *s ∈* {*s*_1_, *s*_2_, …, *s*_*d*_}, we blur the Hi-C image **I**^(*s*)^ ∈ ℝ^*M*×*N*^ and compute two image processing features derived from differential geometry (Haralick, Watson, and Laffey, 1983): the *ridge strength matrix Λ*^(*s*)^ ∈ ℝ^*M*×*N*^ and the *ridge angle matrix* **Θ**^(*s*)^ ∈ ℝ^*M*×*N*^. Suppose that **I**^(*s*)^ is a surface residing in ℝ^3^ where *i*, j defines the position on the image plane and the value **I**^(*s*)^(*i*, j) defines the height. Then, the ridge strength matrix has a large positive value in regions where the surface **I**^(*s*)^ appears to be more ridge-like. A ridge is a point on the surface which has downward facing slopes to its left and right and is relatively flat if you walk in the direction of the ridge. It is possible to quantify how ridge-like any point *i*, j on the Hi-C image is, and likewise, the direction in which the ridge is heading towards (the ridge angle matrix). Both these quantities are computed through the Hessian: the ridge strength is the magnitude of the maximal curvature of the surface **I**^(*s*)^(*i*, j) and the ridge angle is the direction in which there is maximal curvature.

Let *i ∈* {1, …, *M*} be the row index j ∈ {1, …, *N*} be the column index. We construct the 2 × 2 Hessian matrix for each point of the image *i* and j, defined as,

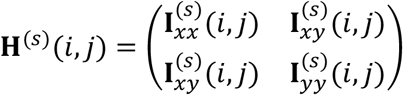

where 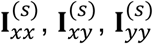 are the outputs of convolving the second order derivative kernels on the blurred Hi-C image I^(*s*)^. Then, we define the ridge strength as a function of the largest eigenvalue of the negated Hessian

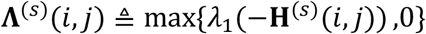

and the ridge angle is the angle of the corresponding eigenvector **v**_1_. Notably, we negate the Hessian matrix such that the largest eigenvalue corresponds to ridges as opposed to valleys, and apply a ReLU (ramp) transformation such that points whose curvature in the direction of the ridge is larger than that in the perpendicular direction is clamped to 0. Practically, we compute both quantities analytically, i.e.,

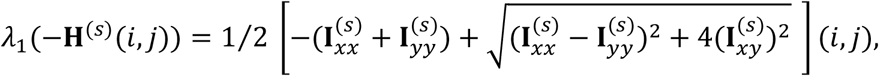

and similarly for the ridge angle,

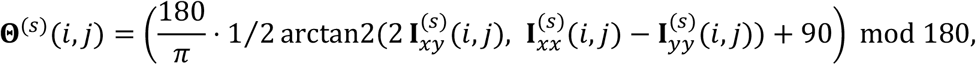

which is the leading eigenvector s_1_ orientation, offset by 90°, converted from radian into degrees and wrapped to [0, 180°). For an example showing both ridge strength and ridge angle matrices, see **Figure S1** ‘Image processing block’.

A subtle yet important aspect of the scale space framework is the derivative kernels, which are convolved with **I**_(*s*)_ to obtain 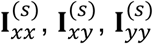 . Let 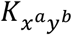 be the discrete kernel taking the *a*-th order *x*-derivative and *b*-th order *y*-derivative. For automatic scale selection (Lindeberg, 1998), it is important that we normalize the output of the discrete derivatives such that the appropriate scale of a ridge appear as a local maximum of *Λ*^(*s*)^ over scale *s*, a property leveraged in *Scale inference*. Define the *γ*-normalized *scale-space derivative kernel*

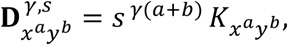

where Lindeberg showed that for a cylindrical Gaussian ridge, *γ* ∈ (0,1.5) must be satisfied to find the correct scale of ridges as the local maxima of *Λ*^(*s*)^ with respect to scale (Lindeberg, 1998). We set the default parameter of *γ* = 0.75 which was suggested for ridges; reducing *γ* = 0.5 is more adapted to find edges, whereas *γ* = 1.0 for blob-like patterns (Lindeberg, 2008). For our application we only need the second order derivatives:

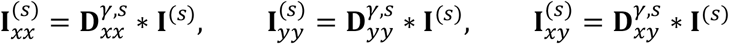

which is computed using function `computeNjetfcn` from *pyscsp* (1.0.0).

### Ridge characterization

#### Ridge detection using ImageJ CurveTrace

MIA-Jet calls the ImageJ *CurveTrace* plugin (0.3.5) on each blurred image **I**^(*s*)^ to obtain ridge coordinates (see **Figure S1** ‘Image processing block’). We use ridge coordinates detected from the CurveTrace to rely on the line linking algorithm proposed by Steger, 1998. The plugin is run with the following parameters: `correct_position`, `estimate_width`, `show_junction_points`, `displayresults`, `method_for_overlap_resolution` is None, `minimum_line_length` is 0, `maximum` is 0, and upper/lower hysteresis thresholds 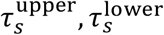 (see *Threshold estimation*). Since the blurred rectangular image **I**^(*s*)^ has number of columns *N* that scales linearly with the chromosome size divided by the resolution (see **Generating an Image from a Hi-C file**), at high Hi-C resolutions, the number of ridges can explode causing ImageJ to crash. To mitigate this, we split the image into evenly sized chunks and call ImageJ on each chunk independently. For each scale *s ∈* {*s*_1_, *s*_2_, …, *s*_*d*_}, we save a results table with ridge coordinates, angle, width, contrast, and asymmetry; tables are then concatenated across chunks (offsetting the ridge coordinates such that the final coordinates are with respect to the entire chromosome) and across scales. Each ridge is tagged with the scale at which it was detected (i.e., we say a ridge is detected *at some scale s*_*i*_ ∈ {*s*_1_, *s*_2_, …, *s*_*d*_} if it was a ridge identified from running CurveTrace on the image **I**^(*s*^_*i*_)).

Suppose that we have detected *K* ridges in total across the entire chromosome numbered from 1,2, … *K*. We introduce the following notation on the *k* th ridge *k ∈* {1,2, …, *K*}:

- The length (the number of points in the ridge): *z*_*k*_
- The row and column ridge coordinates to the Hi-C image as an ordered tuple: ((*r*_1_, *c*_1_), (*r*_2_, *c*_2_), … 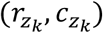
- The scale at which the ridge was detected: 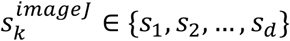

#### Threshold estimation

If the thresholds parameter is not specified, hysteresis thresholds are estimated from image statistics as outlined in the ImageJ *Ridge Detection* plugin (https://imagej.net/plugins/ridge-detection). Let *k*_hi_ and *k*_lo_ be the 75th and 25th percentiles of the non-zero values of I^(*s*)^. First, we convert each scale to widths (in pixels) using the equation from *Choosing the scales*,

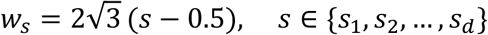

and then define the upper and lower hysteresis thresholds for **I**^(*s*)^

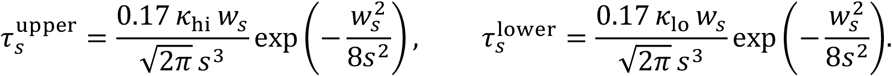

#### Ridge processing

Each ridge obtained from ImageJ CurveTrace is processed as follows. For the *k*th ridge, *k ∈* {1,2, … *K*},

1. Sorting. Ridge coordinates are sorted from smallest to largest distance to the main diagonal. Because the Hi-C image has been rotated 45°, the height of the image corresponds directly to the distance from the main diagonal. Thus it suffices to sort the ridge coordinates according to the row index *r*_*i*_, 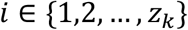. If a ridge *k* is “C-shaped”, which is when the beginning (*r*_1_, *c*_1_) and end coordinates 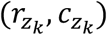 of the ridge are closer to the main diagonal than the coordinate that is furthest from the main diagonal, then no sorting is performed. From hereon we assume that the ridge coordinates ((*r*_1_, *c*_1_), (*r*_2_, *c*_2_), … 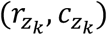 are sorted and refer to the point of the ridge that is closest to the main diagonal (*r*_1_, *c*_1_) as the *root* of a jet.
2. Root enforcement. Jets typically extrude from the main diagonal. To enforce this with some leeway, we discard ridges with root that does not lie within `root_within` from the main diagonal. If root_within ≤ 1 it is a fraction of the window size *H*; if ≥ 1 it is interpreted as the number of bins (default 12 if `exp_type`=‘hic’, 0.5 `exp_type`=‘replihic’). Increasing the `root_within` parameter will capture jets that may not begin extruding from the main diagonal.
3. Strata removal. Ridge points within the `rem_k_strata`-th off-diagonal stratum of the main diagonal are removed (default 1). This parameter is recommended to be set to 1 and its purpose is to filter out false jets where occasionally the main diagonal itself is identified as a jet.
4. Minimum length. Ridges with length *z*_*k*_ ≤ 1 bin are removed, which is too small for a jet.

#### Whitespace trimming

We removed rows and columns ϵ from the Hi-C contact matrix that summed to 0 due to edge-based artefacts from kernel convolution in *Processing Hi-C Data*. After ridge detection (**Ridge characterization**), we add offsets to the ridge coordinates ((*r*_1_, *c*_1_), (*r*_2_, *c*_2_), … 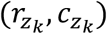 such that the coordinates are corrected. After offsetting, a ridge may “jump” across an unmappable gap.

When such an abrupt coordinate shift occurs, we examine the ridge segment on either side of it: if one side consists of unmappable region (pixel values at or below the 25th percentile of the positive values in the original, unexcluded image), the low-signal segment is discarded and the other segment is kept; otherwise, the ridge is left intact. Ridges reduced to a single point are discarded.

### Combining scale space features with detected ridges

#### Scale inference

This section combines the results of the *Feature Generation* section, scale space feature matrices –ridge strength and ridge angle– with the detected ridge coordinates from the ImageJ CurveTrace plugin, enabling **splitting** and **quantification**. Given the ridge strength matrices {**Λ**^(*s*)^ ∈ ℝ^*M*×*N*^ | *s ∈* {*s*_1_, *s*_2_, …, *s*_*d*_}} and some *k*th ridge *k ∈* {1,2, …, *K*} with coordinates 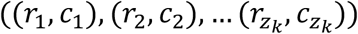, we index the ridge strength matrices at the ridge coordinates using bilinear interpolation, such that we obtain a matrix,

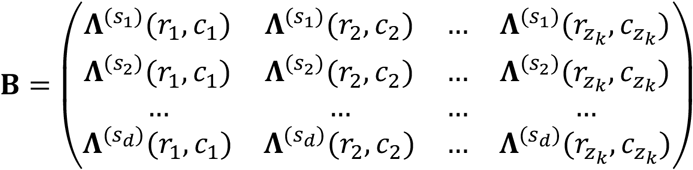

where the columns run across the ridge position and the rows run across the scales (see **Figure S1** ‘Scale inference’). Since we normalized the discrete derivative filters in *Feature Generation*, it suffices that taking the local maxima of the ridge strength with respect to scale gives the appropriate scale of the ridge (Lindeberg 1998). Operationally, this consists of finding the local maxima of each column of **B**, i.e.,

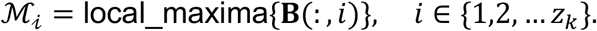

Here, ℳ_*i*_ *⊆* {1,2, …, *d*} will contain the indices of the appropriate scale(s) of the ridge at the *i*th position of the ridge, (*r*_*i*_, *c*_*i*_). We filter out the ridge if there is any column with no local maxima. Notably, there may be more than one scale (|ℳ_*i*_| > 1), which is possible if a single position on the image (*r*_*i*_, *c*_*i*_) has both a large jet and a small jet on top of it. Nevertheless, since this is unlikely to occur in real Hi-C maps, we assign a unique scale to each position (*r*_*i*_, *c*_*i*_) of the ridge by the following method.

First, we combine all the local maxima sets for the entire ridge,

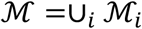

and cluster ℳ using DBSCAN *sklearn*.*cluster*.*DBSCAN* from *scikit-learn* (1.3.2) with fixed parameters `eps` = 2.5, `min_samples` = 2. The aim of DBSCAN clustering is such that when there are multiple local maxima at a position (*r*_*i*_, *c*_*i*_), we select the scales that is most consistent with the entire ridge. Among the non-noise clusters, we select the cluster of scales which intersect with the scale at which the ridge was detected, 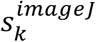, the idea being that since this *k*th ridge was detected from CurveTrace at scale 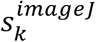, the cluster must contain this scale at least once. If there are multiple clusters that intersect 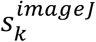, we select the cluster with intersection occurs at the smallest *i*. If no cluster intersects, we select the cluster with scales are closest to 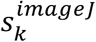 (with ties broken by smallest position *i*). We set `eps` to 2.5 to ensure that a cluster of scales cannot have more than one scale assignment for a column. The output of this step is the final cluster of scales at each position of the ridge,

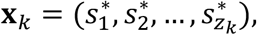

which we refer to as the assigned scales of the ridge. We convert the assigned scales to widths in pixels (*Choosing the scales*) and report the widths in the MIA-Jet output:

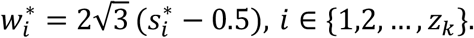

#### Computing Ridge Features

Once we have obtained the assigned scales of the *k*th ridge at each position of the ridge, we can extract the corresponding ridge strength values and angles. For the *k*th ridge, let **r**_*k*_ denote the ridge strength values and **a**_*k*_ the ridge angle values, defined as

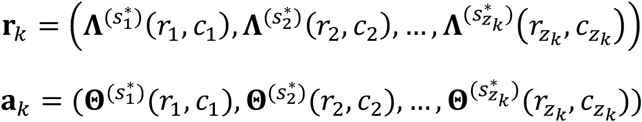

After this step, the *k*th ridge contains information about its strength, angle, and width, denoted by the tuple (**x**_*k*_, **r**_*k*_, **a**_*k*_) associated to each ridge *k ∈* {1,2, … *K*} (see **Figure S1** ‘Scale inference’).

### Generating Compartment Mask

The compartment mask is used in **Splitting** to ensure that identified ridges do not run through A/B compartment boundary intersection points. These cases are often indicative of false jets, or may be two different jets that have been identified as one. The `compartment` parameter (default True for `exp_type`=‘hic’, False for `exp_type`=‘replihic’) controls whether this feature is enabled. This section describes how we compute a compartment mask **I**_compartment_ ∈ {0,1}^*M*×*N*^, which is a matrix of the same dimension as the Hi-C image with 1 where there is a compartment boundary intersection and 0 otherwise.

A/B compartments are defined from the correlation of the O/E map (Lieberman-Aiden et al., 2009). We therefore read in the Hi-C O/E contact matrix *hic-straw* package using `data_type`=‘oe’ and identical `normalization`, and `resolution` as the Hi-C image (*Processing Hi-C Data*). Similar to the Hi-C image, we remove the rows and columns whose sum is 0 and min–max normalize to [0,1], which we denote as **A**_oe_. Then we compute the Pearson correlation matrix

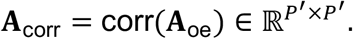

Here, if A_corr_(*i*, j) is close to 1, genomic loci *i* and j are correlated which means that *i* and j should be clustered in the same compartment. Conversely a value of −1 indicates that genomic loci should be in different A/B compartments. We [0,1] min-max normalize **A**_corr_ using `normalize` from *opencv* (4.11.0) such that −1 gets mapped to 0, effectively producing a weighted adjacency matrix of a graph whose edge weights encode compartment affinity.

Finding A/B compartments is then partitioning the nodes of this graph such that intra-connectivity within compartment is maximized and inter-connectivity between compartments are minimized. We find this partition by a principled approach by computing the symmetric-normalized Laplacian

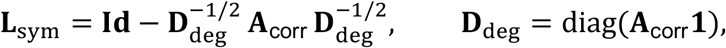

where **Id** is the identity matrix, diag is the diagonal function that creates a square matrix with the given vector placed along its main diagonal, **1** the all-ones vector, and rows/columns of isolated loci are zeroed.

Inspired by the spectral clustering applications, we rely on the fact that the eigenvectors of the Laplacian corresponding to the *t* smallest eigenvalues are the relaxed solutions to finding *t* partitions of genomic loci based on the connectivity of the contact matrix (von Luxburg, 2007). Since there exist A/B compartments and further sub-compartments (Rao et al., 2014), we use both the second and third smallest eigenvalue (discarding the eigenvector corresponding to the smallest eigenvalue since it is the trivial eigenvector). From the framework of spectral clustering, we can expect the eigenvectors to be roughly piecewise constant within a compartment and change sharply at its boundaries, since it solves the relaxed normalized cut problem (Shi and Malik, 2000). We therefore identify A/B compartment boundaries as locations where any of the second and third eigenvectors changes abruptly (i.e. changepoints) by taking the absolute first difference. We binarize the output by thresholding using Otsu’s method (Otsu, 1979) using `threshold_otsu` from *scikit-image* (0.26.0). This is repeated for both eigenvectors and combined with a logical OR to ensure we detect both A/B compartments and sub-compartments, obtaining a vector 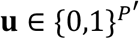 with 1 at the A/B compartment switches.

Finally we compute the outer product matrix 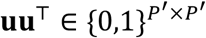 that has a 1 where any two A/B compartment boundaries intersect and 0 otherwise. Intuitively, uu^*⊤*^ has 1 at the corners of the A/B compartment “checkerboard” plaid pattern, which are the points where a compartment boundary along one genomic axis intersects another. The matrix uu^*⊤*^ is rotated 45° and rectangle image is extracted as in **Generating an Image from a Hi-C file**, so that it aligns in dimensions with the Hi-C image **I**. Denote the compartment mask as **I**_compartment_ ∈ {0,1}^*M*×*N*^. This is utilized in the splitting section where ridges that pass through a 1 in **I**_compartment_ are split into two as it is unlikely that a single jet structure extends through A/B compartments. To handle noise, we apply a horizontal dilation kernel to **I**_compartment_ (dimension 1 × 5 if `resolution` < 25 kb 1 × 3 otherwise). As a precaution, we also zero every row within `root_within_comp` bins of the main diagonal so that the roots of any potential true jets are not split too close to the main diagonal (default set equal to `root_within` if `exp_type`=‘hic’).

### Splitting

The hysteresis thresholds supplied to ImageJ CurveTrace may not be accurate or adapted to certain parts of the image but not to smaller local patches (**Ridge characterization**, *Threshold estimation*). Because of this, ridges may extend into noise or through multiple structures. MIA-Jet therefore splits ridges into sub-ridges based on 4 criteria (angle, scale instability, scale decrease, and compartment), ideally separating a ridge if it runs through multiple different structures (e.g. jet, TAD, loops). Each criterion has a parameter which specifies how much of the first few coordinates of a ridge that we protect from splitting (`angle_trim`, `scale_trim`, `scale_dec_trim`, and `comp_trim`). The reasoning behind preventing the first few positions of the ridge from being split is that splitting criteria utilizes (**x**_*k*_, **r**_*k*_, **a**_*k*_) (see *Computing Ridge Features*) to determine where to split, the values of which may contain some noise. To prevent true jets from being split before it ends, we introduce these mechanisms to preserve the root portion of true jets near the diagonal.

Let each trim parameter be represented by *π*_*∗*_ (*π*_angle_trim_, *π*_scale_trim_, *π*_scale_dec_trim_, *π*_comp_trim_). Define *n*_*∗*_ as the protective range of the root of the ridge in number of bins,

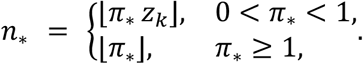

In other words, if a fractional value is given 0 < *π*_*∗*_ < 1, the first *Lπ*_*∗*_*z*_*k*_] positions are protected from splitting (*π*_*∗*_ is interpreted as a fraction of the ridge length *z*_*k*_). If an integer value *π*_*∗*_ ≥ 1, the first *Lπ*_*∗*_] positions are protected (*π*_*∗*_ is interpreted as the number of points of the ridge). Passing None disables the splitting of that criterion entirely. The default for *π*_angle_trim_is 3 if exp_type=‘hic’, 0.5 if exp_type=‘replihic’; *π*_scale_dec_trim_ is 3 if exp_type=‘hic’, None if exp_type=‘replihic’; *π*_scale_trim_, is 0.25 if exp_type=‘hic’, None if exp_type=‘replihic’; *π*_comp_trim_is 3 if exp_type=‘hic’, None if exp_type=‘replihic’. Reducing these values will result in shorter jets since we decrease the protective range near the root. These values were adjusted to find jets in WT DP Thymocytes at 25 kb resolution (Guo et al., 2022) for `hic` mode and Repli-HiC jets at 25 kb resolution (Liu et al., 2024) in `replihic` mode.

Let *i ∈* {1,2, …, *z*_*k*_} denote the ordered position along the ridge, where the point *i* = 1 denotes the point closest to the main diagonal and *i* = *z*_*k*_ denotes the furthest extrusion point. For each of the four criteria we specify the points to split the ridges ℬ_*∗*_:

- **Angle** (`angle_trim`): The ridge positions with angle that is not in the desired angle range [θ_*lb*_, θ_*ub*_] is split, where θ_*lb*_ is the angle lower bound and θ_*ub*_ is the angle upper bound for the jets in degrees, specified by the `angle_range` parameter. Define ℬ_angle_trim_ as the set of ridge positions with angle that is not in the desired angle range [θ_*lb*_, θ_*ub*_]

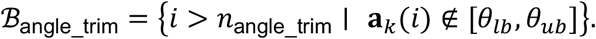
- **Scale instability** (`scale_trim`): The ridge positions where the rolling window standard deviation *ρ*(*i*) of the scale indices exceeds `scale_trim_thresh` = *τ*_sc_ (default 3). The rolling window size is set by the parameter *c* = `scale_trim_window` (default 5). Define *l*_*k*_(*i*) = index(**x**_*k*_(*i*)) for the index of the scale assigned to position *i*, then

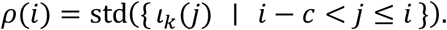

This captures ridges that run through several structures of differing scale, i.e. regions where the assigned scale index varies sharply. Reducing `scale_trim_thresh` will split jets much more often. Define ℬ_scale_trim_ as the set of ridge positions where *ρ*(*i*) exceeds `scale_trim_thresh`,

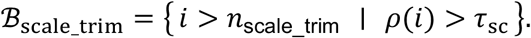
- **Scale decrease** (`scale_dec_trim`): The ridge positions which exhibit a sudden reduction in scale are split. These cases arise when a jet approaches a loop, which registers as a small scale structure. From the ridge root we maintain a running maximum of scales max_*l*≤*i*_index(**x**_*k*_(*l*)); any ridge position which drops from the current maximum by `scale_dec_thresh_trim` = *τ*_dec_ is split (default 10). The ridges are not split if the total number of scales in scale space *d* is less than `scale_dec_thresh_trim`. Define ℬ_scale_dec_trim_ as the set of ridge positions which exhibit a sudden reduction in scale,

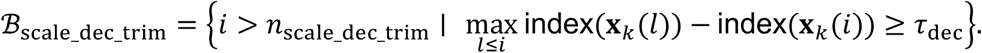

Reducing `scale_dec_thresh_trim` will cause jets to be split more often.
- **Compartment** (`comp_trim`): The ridge positions which go through an A/B compartment boundary intersection point are split (see **Generating Compartment Mask**). Define ℬ_comp_trim_ as the set of ridge positions which go through an A/B compartment boundary intersection point

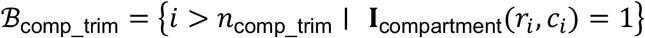

where *r*_*i*_, *c*_*i*_ is the row and column coordinate of the ridge.

Let ℬ = ℬ_angle_trim_ *∪* ℬ_scale_trim_ *∪* ℬ_scale_dec_trim_ *∪* ℬ_comp_trim_. We split the ridge at the positions in ℬ as long as the new split ridge has more than one point (single point segments are discarded).

### Quantification

Quantification assigns metrics to each ridge *k ∈* {1,2, …, *K*} for downstream filtering and ranking. Each ridge carries its assigned scales 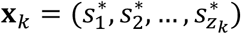, ridge strength values **r**_*k*_ and angle values **a**_*k*_ (see *Computing Ridge Features*). Let [θ_*lb*_, θ_*ub*_] be the `angle_range`. For each *k*th ridge, *k ∈* {1,2, …, *K*} we compute jet saliency and consistency metrics (used for ranking jets), and turbulence metrics (used for filtering).

#### Jet saliency

Jet saliency is the weighted sum of ridge strength values 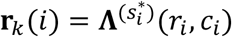,

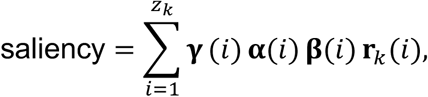

where **r**_*k*_(*i*) is the ridge strength at position *i* evaluated at its assigned scale 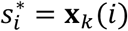 (*Scale inference*), and weights *γ*(*i*), **β**(*i*), and *α*(*i*) encode desirable jet characteristics, such as whether the jet angle is in the user-specified range `angle_range`:

- **Angle condition** at the assigned scale:

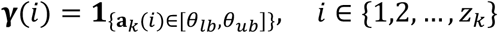

captures whether the ridge is heading in a direction that the user specifies via `angle_range` parameter.
- **Angle condition** across scales:

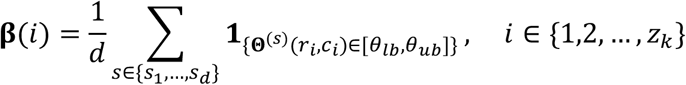

measures how consistently the angle is in the desired `angle_range` across all scales. It is the fraction of scales at which the angle at (*r*_*i*_, *c*_*i*_) stays within [θ_*lb*_, θ_*ub*_]. A true jet typically has angles that remain consistent across scales, whereas false jets from loops or TADs exhibit significant variation in angle across the scales. The `ang_frac` parameter (default True if `exp_type`=‘hic’, False if `exp_type`’replihic’) controls whether **β**(*i*) is taken into consideration in jet saliency and consistency metrics. It is not taken into account by setting **β**(*i*) = 1 for all *i ∈* {1,2, …, *z*_*k*_} causing no meaningful contribution to the jet saliency as the weight is all 1. Setting `ang_frac` to False can find jets that are more variable in curvature.
- **Non-decreasing scale** *α*(*i*). Diffuse chromatin jets broaden gradually with distance from the main diagonal (Guo et al., 2022), which in scale space corresponds to a non-decreasing assigned scale along the ridge from the root to the end. We capture this with *α*(*i*), a boolean sequence computed from the assigned scale indices index(**x**_*k*_(*i*)) ordered from the root position *i* = 1. Let *α*(1) = 1, and for *i* ≥ 2

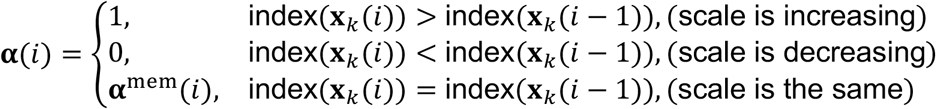

where the memory value *α*^mem^(*i*) equals the most recent strict change (1 after an increase, 0 after a decrease). The memory value helps with robustness to noise. For ridges with an increasing scale progression (i.e., a jet that broadens as it extrudes, see **Figure S1** Jet 1) *α*(*i*) is largely populated with 1s. The `adj_nondec` parameter (default True if `exp_type`=‘hic’, False if `exp_type`’replihic’) controls whether *α*(*i*) is taken into consideration in jet saliency and consistency metrics. Similar to **β**(*i*), it is not taken into account by setting *α*(*i*) = 1 for all *i ∈* {1,2, …, *z*_*k*_}. Setting `adj_nondec` to False can help finding jets that do not necessarily broaden as it extrudes (**Figure 3a**). The jet saliency measure was based on *ridge saliency* (Lindeberg, 1998, equation 58):

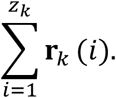

Our modification to finding jets is in weighting this measure such that we encourage jets with user-defined properties, namely the angle in which the jet is extruding (`angle_range`) and whether or not we want to capture diffuse chromatin jets. This notion of weighting by desirable properties is extended in consistency metrics.

#### Consistency metrics

Consistency metrics are designed to directly quantify jets with a non-decreasing scale without necessarily taking into account ridge strength. This is useful at picking up jets with relatively weak ridge strength but which nevertheless look like a jet (see jet chr3:63-64.8Mb **Figure S1** ‘Quantification and Filtering’).

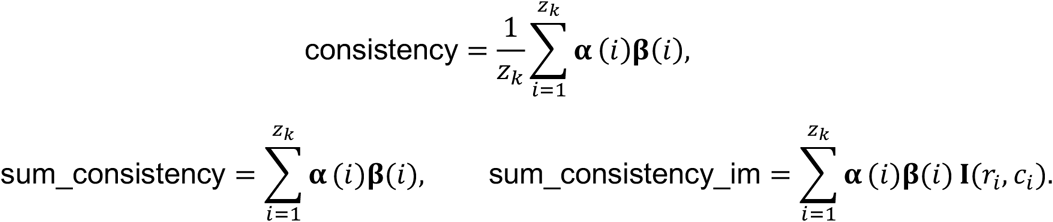

Consistency measures the fraction of the ridge with a non-decreasing scale (*α*), while sum consistency counts the number of ridge points with a non-decreasing scale. Sum consistency is the final ranking metric for jets in Hi-C data (**Figure S1** ‘Quantification and filtering’). Sum consistency im additionally weights by image intensity I(*r*_*i*_, *c*_*i*_) to downweight jets found in sparse, low-signal regions and is the final ranking metric for jets in Repli-HiC data, which exhibits sparse regions.

#### Turbulence metrics

Turbulence metrics capture the variation of signal in either the ridge strength or the angle. False jets often register in regions where the angle is inconsistent or ridge strength has large variation. Turbulence metrics are used in **Filtering** to filter out jets. Let CV(*⋅*) denote the coefficient of variation std/(mean). The angle_turbulence is the CV of the angle over all scales and positions of the ridge,

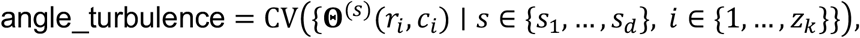

and the ridge_strength_turbulence is the CV of the ridge strength along the assigned scales,

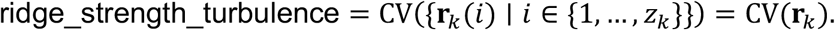

#### Other metrics

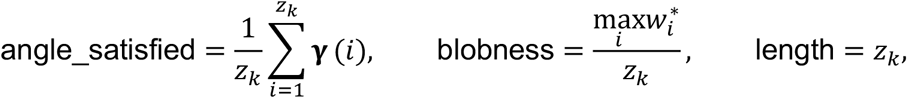

where 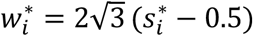 is the width at the assigned scale (*Scale inference*). Angle satisfied is the fraction of points with angle in [θ_*lb*_, θ_*ub*_], and blobness is the ratio of the maximum scale width to the ridge length (used to reject blob-like detections in **Filtering**).

### Computing p-values

#### Whitened Hi-C Image

We compute the *p*-value from the local enrichment of a ridge relative to its surrounding region using another variant of a Hi-C map: I_obs_ ∈ ℝ^*M*×*N*^. The pixel values used for this test are taken from the “observed” data type contact map. Otherwise, the resulting image I_obs_ ∈ ℝ^*M*×*N*^ is generated by the same processing as the Hi-C image I (rotation by 45° and extracting a rectangle) in **Generating an Image from a Hi-C file**.

When `compartment` is True, we compute an additional *p*-value on a Hi-C image with the strength of A/B compartments and TADs reduced. The motivation is that such structures can register as significant local enrichment, so that a small *p*-value reflects a compartment or a TAD rather than a true jet (**Figure S1** ‘P-Value Computation’).

To generate the whitened Hi-C image, we read in the O/E map with indices ϵ removed, followed by log transformation and min–max normalization to [0,1] giving the square matrix 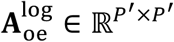. Let center(*⋅*) denote double centering (subtracting the row means and column means, then adding back the global mean). Define the covariance matrix,

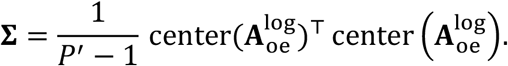

The A/B compartment eigenvector is the leading eigenvector of the Pearson correlation matrix of the contact profiles. Here, Σ is constructed from the same centered cross-product, differing only in that it is symmetrically double-centered rather than rescaled to unit variance. Thus, both the correlation and covariance matrix are dominated by the same large-scale structure A/B compartments. We use the covariance matrix as the whitening operation requires a covariance matrix.

To whiten the image, we first eigendecompose the covariance matrix Σ = Σ diag(***λ***) V^⊤^ where 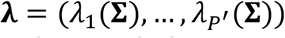 are the eigenvalues and the columns of 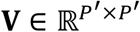 are the corresponding orthogonal eigenvectors. Define the whitening matrix (Bell and Sejnowski, 1997)

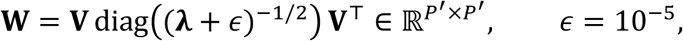

where *⋅*^−1/2^ is applied elementwise.

Following the works of Bell and Sejnowski, the whitened matrix is then,

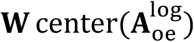

which is processed by adding the absolute value of its minimum entry to ensure that values are non-negative. The whitened matrix is rotated 45° and we extract a rectangular region as in **Generating an Image from a Hi-C file** to give **I**_white_ ∈ ℝ^*M*×*N*^.

The whitening operation flattens the eigenvalue spectra of the Hi-C contact matrix. Because the A/B compartments are the dominant modes of the Hi-C contact matrix (Lieberman-Aiden et al., 2009), the whitening operation effectively reduces the strength of the lower rank, A/B compartment structure. Applying the enrichment test on **I**_white_ can measure local enrichment that is more sensitive to diagonally dominant structures than just **I**_obs_. **Figure S1** shows an example region where a false jet (jet 2) has less enrichment on the whitened image compared to the observed image.

### Statistical Test

MIA-Jet assigns each ridge a p-value measuring the local enrichment of the ridge relative to its flanking regions (**Figure 1d**, “Statistical test”).

Consider the *k*th ridge *k ∈* {1,2, …, *K*} with coordinates ((*r*_1_, *c*_1_), (*r*_2_, *c*_2_), …, 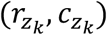 and widths 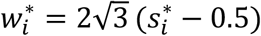For each ridge point (*ri, ci*), let 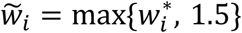 be the box width (in pixels) and define three congruent 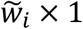 boxes: a **center** box *C*_*i*_ centered at (*r*_*i*_, *c*_*i*_) with its long axis oriented across the ridge (the normal of angle **a**_*k*_(*i*)), and **left** and **right** boxes ℒ_*i*_, ℛ_*i*_ obtained by translating *C*_*i*_ by 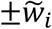 along that axis, so that they tile the adjacent flanks (**Figure 1d**, “Statistical test”).

For a given Hi-C image, let 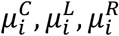 be the mean pixel values of that image within the respective boxes *C*_*i*_, ℒ_*i*_, ℛ_*i*_, and define the flanking background by the geometric mean

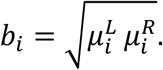

Let 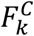and 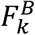 be the empirical CDFs of 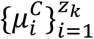 and 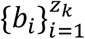. We perform a one-sided two-sample Kolmogorov–Smirnov test with statistic

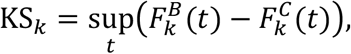

whose null hypothesis is that the center is no larger than the background and whose alternative is that the center is larger using `ks_2samp` from *scipy* (1.16.1). We carry out this procedure on the observed image **I**_obs_ to obtain the p-value *p*_*k*_ (`p-val`) and, in compartment mode, repeat it on also **I**_white_ to obtain 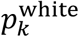 (`p-val_white`).

### Filtering

Ridges are filtered by thresholds. Each threshold is optional and specifying it directly will override the default assigned by `exp_type`. A value of `None` disables it and ‘–’ indicates that the filter is always applied and cannot be explicitly set by the user.

| Filter Parameter | Description | Discard rule | Default `exp_type`='hic' \| `exp_type`='replihic' |
| --- | --- | --- | --- |
| *length* | Removes ridges smaller than *length* points | `metric < threshold` | 5 \| 5 |
| *angle_turbulence* | Removes jets with angles that significantly vary | `metric ≥ threshold` | 0.325 \| None |
| *ridge_strength_turbulence* | Removes jets with ridge strength that significantly vary | `metric ≥ threshold` | 1.0 \| None |
| *blobness* | Remove blob-like jets | `metric ≥ threshold` | 2.0 \| 2.0 |
| *consistency* | Removes jets that do not have a broadening width | `metric < threshold` | 0.6 \| None |
| *sum_consistency* | Ranking metric for Hi-C jets | `metric < threshold` | None \| None |

| Filter Parameter | Description | Discard rule | Default *exp_type*='hic' \| *exp_type*='replihic' |
| --- | --- | --- | --- |
| *sum_consistency_im* | Ranking metric for Repli-HiC jets (also applicable for other sparse contact maps) | metric < Yen threshold (automatic) | False \| True |
| *angle_satisfied* | Remove jets with angles not in the desired *angle_range* | metric < threshold | 0.3 \| 0.3 |
| *saliency* | Remove jets that do not have significant local enrichment | metric < Triangle threshold (automatic) | — |

The Yen and Triangle thresholds (Yen et al.,, 1995; Zack et al.,, 1977) are computed automatically from the distribution of the metric across all ridges using `threshold_yen`, `threshold_triangle` from *scikit-image* (0.26.0).

### Overlap removal

Since ridge detection runs at multiple scales, several ridges may annotate at the same location. We iterate over all ridges, identify overlapping groups, and keep a single ridge per overlapping group. Any two ridges are deemed to intersect if they overlap in their coordinates and the intersection over union exceeds 0.1. Among overlapping jets, we select the jet that has the maximal or minimal `resolve_conflict` statistic, which is minimized for `p-val`, `p-val_white`, `blobness`, and maximized for `length`, `saliency`, `sum_consistency`. The default `combined` is the average of the −log_10_ *p*-value and the saliency.

### Multiple-test correction and significance thresholding

The Benjamini–Hochberg procedure using `multipletests` with `method =` *‘*`fdr_bh`*’* from *statsmodels* (0.14.5) is applied separately to {*p*_*k*_}_*k∈*{1,2,…*K*}_, giving corrected values {*q*_*k*_}_*k∈*{1,2,…*K*}_ and, in compartment mode, to 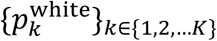, giving 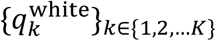. Ridges are retained if *q*_*k*_ ≤ `q_val` (default 0.1 if `exp_type`=‘hic’, 0.2 if `exp_type`=‘replihic’). In compartment mode, an additional filter is applied: 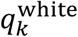 ≤ `q_val_white` (default 0.95).

### Outputs

For each run, three results are produced: (1) all ridges after processing (*Ridge processing*), (2) filtered ridges (*Filtering* and *Overlap removal*), and significant ridges (*Multiple-test correction and significance thresholding*), in a subfolder named for the significance cutoffs. Each result comprises:

- **Summary table** (one row per ridge): unique_id, which is the unique ID of the jet, chromosome, start, and end, which are the genomic coordinates of the jet root *r*_1_, *c*_1_ (closest point to the main diagonal),s_imagej, the scale at which the ridge was detected 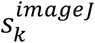, length, which is the number of points in the jet, dist_diag, which is the distance of jet root *r*_1_, *c*_1_ to the main diagonal, all quantification metrics, *p* and *q*-values.
- **Expanded table** (multiple rows per ridge): unique_id, which is the unique ID of the jet, chromosome, x (bp) and y (bp) which are the genomic coordinates of ridge positions, x (pixels) and y (pixels) which are the coordinates of the ridge positions with respect to the binned Hi-C image (*r*_1_, *c*_1_), … 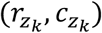, assigned scales **x**_*k*_, jet angles **a**_*k*_, ridge strength **r**_*k*_, width 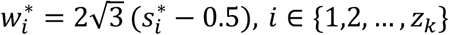, and input, which are the Hi-C image values along the jet. In addition, the asymmetry and contrast are carried over from the CurveTrace plugin.
- **Visualization** .**bedpe file**: .bedpe file for visualization in Hi-C visualization software (e.g. Juicebox or HiGlass).
- **Output plot**: the Hi-C image with detected ridges overlaid
- **Diagnostic plots**: The –diagnostic_plots parameter will generate plots showing jets after each operation in splitting, filtering, and p-value filtering.

Ridges are linked across tables by the `unique_id` column. MIA-Jet runs each chromosome independently, so results are submitted in parallel with script *submit_all*.*py* with the parameters specified in *submit_all_config*.*yaml*. Results across chromosomes can be combined with *submit_combine_chrom_results*.*py*.

### Influence of Contact Density

MIA-Jet can work with a range of sparsity levels for 3C data. In Hi-C mode, it is recommended that there are at least 480 million number of intra-chromosomal pairwise interactions. If the Hi-C data has significant sections of unmappable regions, setting `sum_consistency_im` to `True` may help remove jets identified at unmappable regions. In RepliHi-C mode, it is recommended that there are at least as or more than 32 million number of intra-chromosomal pairwise interactions.

### Analysis of Results

The datasets used to analyze the MIA-Jet results are listed in **Supplementary Table S1** ‘Analysis Data’.

#### Generating simulated data

We generated simulated Hi-C data using 3DPolyS-LE (Gitchev et al., 2022): 7 chromosomes containing jets and 2 chromosomes containing TADs as false jets. Here is an outline of how individual chromosomes were generated and the rationale for the choice of parameters:

1. Unbounded jets with no CTCF.
  1. Chromosome size: 40 Mb
  2. Number of loading sites: 20, evenly spaced across the chromosome. We randomly grouped loading sites into 10 “narrow” and 10 “broad” where the narrow loading site is 50 bp and broad is 10 kb.
  3. No CTCF boundary sites
  4. Generated chromosomes:
    1. `test1a_fast` (**Figure 2a**): Generated a chromosome with *3DPolyS-LE jet parameters* using the 10”narrow” loading sites only and fast cohesin extrusion speed *km* = 1.5 × 10^−2^.
    2. `test1a_slow`: Generated a chromosome with *3DPolyS-LE jet parameters* using the 10 “broad” loading sites only and slow cohesin extrusion speed *km* = 5 × 10^−4^; for broad loading sites the cohesin speed had to be reduced to observe jets.
2. Bounded jets with convergent CTCF
  1. Chromosome size: 40 Mb
  2. Number of loading sites: 20, evenly spaced across the chromosome. We randomly grouped loading sites into 5 “short”, 5 “medium”, 5 “long”, and 5 “super long”. The loading site size was fixed to 50 bp.
  3. CTCF boundary sites: For each group, we placed convergent CTCF boundaries of equal distance from the loading site, where the distance is randomly sampled from a uniform distribution [50 kb, 250 kb] for short, [250 kb, 500 kb] for medium, [500 kb, 1 Mb] for long, and [1 Mb, 2.25 Mb] for super long. We made sure that no CTCF boundary between two jets were overlapping by placing a buffer of 750 kb between any CTCF motifs between two jets. The CTCF boundary impermeability strength was 1.
  4. Generated chromosomes:
    1. `test1b`: Generated a chromosome with *3DPolyS-LE jet parameters* and fast cohesin extrusion speed *km* = 1.5 × 10^−2^.
3. Curved jets (small jets)
  1. Chromosome size: 40 Mb
  2. Number of loading sites: 21, evenly spaced across the chromosome. We randomly grouped loading sites into 7 “left”, 7 “right” 7 “straight”. The loading site size was fixed to 50 bp
  3. CTCF boundary sites: For “left” jets, we placed a forward orientation CTCF motif upstream of the loading site at a random distance drawn from a uniform distribution. Similarly, for “right” jets we placed a reverse orientation CTCF motif downstream of the loading site at a random distance. For “straight” jets we placed no CTCF motifs on either side. The CTCF boundary impermeability strength was 1.
  4. Generated chromosomes:
    1. `test1c`: Generated a chromosome with *3DPolyS-LE jet parameters* and fast cohesin extrusion speed *km* = 1.5 × 10^−2^.
4. Curved jets (large jets). Identical settings to curved jets (3) but with 15 loading sites instead of 21 with 5 “left”, 5 “right” and 5 “straight” jets. This creates the potential for larger jets. The motivation was to vary both the size and curvature of jets to a greater extent.
  1. Generated chromosomes:
    1. `test1cii` (**Figure S2a**): Generated a chromosome with *3DPolyS-LE jet parameters* and fast cohesin extrusion speed *km* = 1.5 × 10^−2^.
5. Unbounded jets with no CTCF (Poisson process). Since the chromosomes above placed loading sites on an evenly spaced grid, we decided to introduce more potential variation in jet location and sizes by a Poisson process.
  1. Chromosome size: 20 Mb
  2. Loading sites: We first modeled boundaries as inter-arrival times with exponential distribution mean 500 kb. For each boundary, we performed an independent Bernoulli experiment with probability 0.12 for whether to place a jet or not. For each jet, we placed a loading site at the midpoint of the boundary locations with a loading site size that is randomly sampled from a uniform distribution [1 bp, 100 bp]. Notably, the boundaries were simply used to assign the loading sites and no CTCF boundary elements were placed.
  3. No CTCF boundary sites.
  4. Generated chromosomes:
    1. `chromsim1-fast-unbounded`: Generated a chromosome with *3DPolyS-LE jet parameters* and fast cohesin extrusion speed *km* = 1.5 × 10^−2^.
6. Bounded jets with convergent CTCF (Poisson process).
  1. Chromosome size: 20 Mb
  2. Loading sites: We first modeled boundaries as inter-arrival times with exponential distribution mean 500 kb. For each boundary, we performed an independent Bernoulli experiment with probability 0.12 for whether to place a jet or not. For each jet, we placed a loading site at the midpoint of the boundary locations with a loading site size that is randomly sampled from a uniform distribution [1 bp, 100 bp].
  3. CTCF boundary sites: We assigned for each jet to have only one CTCF motif upstream of the loading site (with probability 0.25), downstream of the loading site (with probability 0.25) or two CTCF motifs upstream and downstream (with probability 0.5). The CTCF boundary impermeability strength was set to 0.8 to promote a more “smooth” curvature once the extruding jet encounters the CTCF.
  4. Generated chromosomes:
    1. `chromsim1-fast-bounded`: Generated a chromosome with *3DPolyS-LE jet parameters* and fast cohesin extrusion speed *km* = 1.5 × 10^−2^.
7. TADs
  1. Chromosome size: 40 Mb
  2. Number of loading sites: 0. Although we placed no specific loading sites, we ensured there was basal loading on the DNA strand.
  3. Number of convergent CTCF motif pairs: 20. We randomly grouped each pair of convergent CTCF motif into 5 “short”, 5 “medium”, 5 “long” and 5 “super long”. For each group, we placed convergent CTCF boundaries of distance between each other that is randomly sampled from a uniform distribution [100 kb, 500 kb] for short, [500 kb, 1 Mb] for medium, [1 Mb, 2 Mb] for long, and [2 Mb, 4.5 Mb] for super long. We made sure that no CTCF boundary between two pairs were overlapping by placing a buffer of 750 kb between any two pairs of CTCF motifs. Although CTCF motifs had orientation, we made the CTCF motif block cohesin extruding on both sides as this created the strongest TAD patterns. The CTCF boundary impermeability strength was 1.
  4. Generated chromosomes:
    1. `test1e`: Generated a chromosome with *3DPolyS-LE TAD parameters* and *z*_*loop* = *True*.
    2. `test1eii` (**Figure S2b**): Generated a chromosome with *3DPolyS-LE TAD parameters* and *z*_*loop* = *False*.

#### Simulation parameters

- **3DPolyS-LE jet parameters**: We optimized parameters to observe jets; all parameters are in the recommended ranges and use the same formulas presented in Gitchev et al., 2022: *NChain* = (*chromosome*_*size*)/(2000), *L* = *round*((*NChain*/2)^1/3^), *kb* = 6 × 10^−6^ (60% cohesin bound to chromatin), *ku* = 2 × 10^−6^, *basal*_*loading*_*factor* = 0, and *Nlef* = (1/*cohesn*_*density*)(2*NChain*)(*kb* + 2*ku*)/*ku*, where *cohesin*_*density* = 50 corresponding to 1 bound cohesin per 50 kb.
- **3DPolyS-LE TAD parameters**: Polymer simulation of TADs had existing example parameters (Gitchev et al., 2022), which we used directly: *NChain* = (*chromosome*_*size*)/(2000), *L* = *round*((*NChain*/2)^1/3^), *kb* = 2.8 × 10^−6^, *ku* = 2 × 10^−6^, and *Nlef* = (1/*cohesn*_*density*)(2*NChain*)(*kb* + 2*ku*)/*ku*, where *cohesin*_*density* = 60 corresponding to 1 bound cohesin per 60 kb. Since we did not explicitly specify loading sites, we set *basal*_*loading*_*factor* = 0.1.

#### Running Fun (Liu et al.,, 2024) on simulated data

We ran the Fun pipeline (v1.0.1) on simulated data at 25 kb resolution. `calculate-son-score` was run with parameters `use_mean True`, `--coverage_ratio 0`, `--padding_width 2`. The `--offset` parameter was set to 5 times the Hi-C resolution and `--ext_length` as (*Window*_*size* − *Window*_*size*/6)/2 ≈ 2.1 Mb where Z*indo*Z_*size* = 5 Mb. `find-fontains` was run with the same parameter values as `calculate-son-score` for `--ext_length`, `--offset`, `--coverage_ratio`. Other parameters were specified as the following: *–*`-half_width 2`, `--extension_pixels 10,100,5`, `--interval_length` as 5 times the Hi-C resolution, `--max_merge_distance` as 5 times the Hi-C resolution, *--p_value 0*.*05*, *--signal_noise_background 1*.*25*. Parameter variation for resolution was run at 10kb, 25kb, 50kb, and 100kb keeping other parameters as described above. Parameter variation for extension length `ext_length` was run with window sizes 1 Mb, 2.5 Mb, 5 Mb, 7.5 Mb, and 10 Mb corresponding to `ext_length` 417kb, 1.04 Mb, 2.1 Mb, 3.1 Mb, and 4.2 Mb keeping other parameters as described above.

#### Running Fun (Liu et al.,, 2024) on mouse DP Thymocytes and human K562 cell-line

We ran the Fun pipeline (v1.0.1) on DP Thymocytes and K562 cell-line ICE normalized mcool files at 50kb resolution. `calculate-son-score` was run with parameters `--norm weight`, `use_mean True`, `--coverage_ratio 0`, `--padding_width 2`. The `--offset` parameter was set to 5 times the Hi-C resolution and `--ext_length` as (*Window*_*size* − *Window*_*size*/6)/2 = 2500000 where *Window*_*size* = 6 Mb. `find-fontains` was run with the same parameter values as `calculate-son-score` for `--ext_length`, `--norm`, `--offset`, `--coverage_ratio`. Other parameters were specified as the following: *–*`-half_width 2`, `--extension_pixels 10,100,5`, `--interval_length` as 5 times the Hi-C resolution, `--max_merge_distance` as 5 times the Hi-C resolution, *--p_value 0*.*05*, *--signal_noise_background 1*.*3*.

#### Definition of K562 Type I and Type II fountains

We downloaded Type I and Type II fountains identified in K562 Repli-HiC data (Liu et al.,, 2024). As described in Liu et al., 2024, K562 Type I fountains are relatively small jets which correspond to replication initiation zones. It is identified from K562 Repli-HiC data using combined Fun results at 5kb, 10kb, and 20kb and are further filtered to overlap with replication initiation zones or a positive origin efficiency metric score from OKseqHMM. Meanwhile, Type II fountains are larger and correspond to replication termination zones. They are identified using Fun results at lower resolutions 50kb, 100kb, and 200kb and filtered to overlap with replication termination zones or a negative origin efficiency metric score. See Liu et al.,, 2024 for more details.

#### Running Fontanka (Galitsyna et al., 2026) on simulated data

We ran Fontanka (v0.1) on simulated data at 25kb resolution. The simulation Hi-C files were ICE normalized using *cooler*.*balance_cooler*. `slice-windows` was run with window size `-W 500000`, `--view` set to the chrom.sizes file and `--expected` with expected cis contact frequencies generated from *cooltools*.*expected_cis*. We utilized the binary fountain mask by calling `apply-binary-fountain-mask` with *-A 0*.*3491* (20 degrees) and the same parameters otherwise as `slice-windows` for `-W` and `--view`. According to example code (Galitsyna et al., 2026), we applied three different thresholding schemes and selected the one which maximized the F1 score: we first dropped missing entries from the results table and applied thresholding to the “FS_peaks” column with the maximum of the Li threshold (*skimage*.*filters*.*threshold_li*) and 0 (to require positive correlation). Parameter variation for resolution was run at 10kb, 25kb, 50kb, and 100kb keeping other parameters as described above. Parameter variation for window size `-W` was run at 100kb, 500kb, 1Mb, 2Mb, and 3Mb.

#### Running Fontanka (Galitsyna et al., 2026) on mouse DP Thymocytes and human K562 cell-line

We ran Fontanka (v0.1) on DP Thymocytes and K562 cell-line at 50kb resolution. The Hi-C files were ICE normalized using *cooler*.*balance_cooler*. `slice-windows` was run with window size `-W 2000000`, `--view` set to *bioframe*.*make_chromarms* with centromeres from *bioframe*.*fetch_centromeres* and `--expected` with expected cis contact frequencies generated from *cooltools*.*expected_cis*. We utilized the binary fountain mask by calling `apply-binary-fountain-mask` with *-A 0*.*3491* (20 degrees) and the same parameters otherwise as `slice-windows` for `-W` and `--view`. Since K562 RepliHiC data largely consisted of jets with few other structures (TADs or loops), we applied the thresholding applied scheme in *examples/00_fontanka_zebrafish*.*ipynb* (Galitsyna et al., 2026): we dropped missing entries from the results table and applied thresholding to the “FS_peaks” column with the maximum of the Li threshold (*skimage*.*filters*.*threshold_li*) and 0 (to require positive correlation). For DP Thymocytes, which contains TADs and loops, we applied a stringent thresholding method presented in “examples/02_fontanka_xenopus.ipynb” (Galitsyna et al., 2026), which applied the same thresholds as described above, followed by additional filtering, keeping jets above the 90th percentile of “FS_peaks” values and jets below the 75th percentile of “Scharr_box” values.

#### Running MIA-Jet on simulated data

We ran MIA-Jet (v2.0.3) on simulated data at 25kb resolution with `--exp_type` *“*`hic`*”*, `--compartment` *“*`False`*”*, `--normalization` *“*`NONE`*”*`--window_size` *“*`5000000`*”*, `--root_within` *“*`5`*”*, `--q_val` *“**0*.*25**”*, and `--angle_range 60 120` to compute the performance scores (**Figure 2b, c**). We then ran MIA-Jet with the same parameters except extending the angle range `--angle_range 45 135` to capture a greater deviation in jet curvature and classify them as “curved” or “straight”. Jets were classified as “straight” if more than 50% of its points had angle in the range [70°, 110°], and otherwise as “curved” (**Figure 2a, Figure S2a, b**, last row).

#### Running MIA-Jet on all cell-lines

We ran MIA-Jet (v2.0.3) with parameters specified in **Supplementary Table S1** ‘Analysis Parameters’. All reported results were submitted in parallel using a python script *submit_all*.*py* and combined across chromosomes with *submit_combine_chrom_results*.*py*. We used the output that is the significance q-val thresholded.

#### Generating downsampled WT DP Thymocytes

Downsampled paired-end FASTQ files were generated for WT DP Thymocytes at 75%, 50%, 25% and 10% using `sample` from *seqtk* (1.3). FASTQ files were aligned to the mm10 genome and converted to Hi-C contact matrices using the *Juicer* pipeline (v2.0) with MboI restriction-site file.

#### Computing overlaps between two sets of jets

Given two sets of jets A and B, we first stratify jets by chromosome such that the intersection of jets for each chromosome independently is computed separately and summed to get the number of intersections for a genome-wide intersection count. Since jets are two-dimensional objects which can intersect with multiple other jets, we compute the intersection over union IoU between A and B in a “left join” fashion: every jet in A records if it has any intersection(s), potentially multiple, with the jets in B. Similarly, we then compute the IoU between A and B in a “right join” fashion. We then construct a bipartite graph where the nodes are jets in A and B. A directed edge is constructed from A to B or B to A if it has been detected from the “left join” or “right join”, respectively, with edge weight given by the IoU values. Here, we seek an optimal 1-1 matching between the jets in A and B by maximizing the total matched IoU using the `max_weight_matching` function from *networkx* (3.3). We expand the coordinates of each jet by a radius of 3 bins using *shapely* (2.1.1) before computing the intersections, where the bin size is the resolution that the jets were called at.

#### Variable length aggregate jet maps

Since jets can have different sizes, we generated aggregate maps where the region is defined by the extrusion point of each jet. We used the extrusion point as the “start” and “end” of the region and rescaled the matrix to be 100×100 using `resize` from *opencv* (4.12.0). Each region matrix was then averaged element-wise to yield the aggregate jet maps.

#### Fixed length aggregate jet maps

We generated aggregate maps where the region is a fixed size extending from the midpoint of the “start” and “end” coordinates. Each region matrix was then averaged element-wise to yield the aggregate jet maps.

#### Signal enrichment at jet locations

Given ChIP-seq, insulation score, compartment score, and repli-seq tracks, we quantified the enrichment of jet locations with respect to these signals. For each jet, we computed the midpoint of the “start” and “end” coordinates, which is the projection of the closest position of the jet to the main diagonal. We then extend the interval by ± 2 bins away from the jet midpoint, where the bin size is the resolution that the jets were called at. These intervals are used to extract the max signal of ChIP-seq, insulation scores, compartment scores, and repli-seq data using the `stats` function from *pyBedGraph* (0.5.43). Any two sets of signal enrichment values are tested for significance by the two-sided Wilcoxon rank-sum test using `ranksums` from *scipy* (1.16.1).

#### Generating random control

Given a set of jets which constitutes the “Observed” data, we generate a random control by the `random` function from *pybedtools* (0.12.0) with the length `l` parameter as the median of the MIA-Jet jets and the number `n` as the number of jets of the set. The random generation was stratified by chromosome and combined.

#### ECDF closest distance to replication zones

For each jet, the closest distance to either the “start” or “end” of a replication zone is computed by `closest` from *pybedtools* (0.12.0) with `d=True` to record the genomic distance and `t=`*“*`first`*”* to handle ties. Each resulting distance is treated as a realization of a random variable, and we estimate its underlying distribution using the empirical cumulative distribution function (ECDF). To test for significant difference between two distributions, we perform the two-sided Kolmogorov–Smirnov test using `ks_2samp` from *scipy* (1.16.1).

#### Number of NIPBL peaks per jet

Given a set of jets where “start” and “end” interval defines the jet loading site, we extend each interval by ±1 bin, where the bin size is the resolution that the jets were called at (25kb for Hi-C GM12878 and K562 results) using the `slop` function from *pybedtools* (0.12.0). The NIPBL peaks are then intersected with the extended jet intervals using `intersect` from *pybedtools* (0.12.0) with `u=True` such that we count distinct peaks only once if it overlaps multiple jets. We then divide the number of total intersected peaks with the number of jets called to get the number of peaks per jet. We repeat this procedure for 3 distinct sets of jets: jets specific to K562, jets specific to GM12878, and jets common to K562 and GM12878, which are derived from *Computing overlaps between two sets of jets*.

#### HCT116 fraction overlap with replication zones

Replication initiation zones (IZs) are genomic intervals that can be “early”, “earlymid”, “latemid”, “late”. We extend each IZ interval by ± 1 bin, where bin size is the resolution that the jets were called at (10kb for HCT116) using the `slop` function from *pybedtools* (0.12.0). Given a set of jets, we compute the midpoint of “start” and “end”, which is the projection of the closest position of the jet to the main diagonal. We then extend the interval by ± 1 bin away from the jet midpoint, where the bin size is the resolution that the jets were called at (10kb for HCT116). A left outer join between the jet intervals and the IZs using `intersect` from *pybedtools* (0.12.0) with `loj=True` is performed, such that every jet is kept whether or not it overlaps an IZ. Each jet is then assigned to one category: either a replication timing category (“early”, “earlymid”, “latemid”, “late”) or “none” if it overlaps with no IZ. If a jet is overlapping multiple IZs, the highest priority is given to the category with the largest fraction of overlap with the jet. This procedure is repeated on a matched set of randomly sampled jets (*Generating Random Control*).

#### Jet angle CTCF motif boxplot

We aimed to find the jet angles at CTCF motif where there is likely CTCF protein binding. To do this, for each CTCF motif interval with “start” and “end”, we extracted the max CTCF ChIP-seq signal intensity value using the `stats` function from *pyBedGraph* (0.5.43) and filtered out CTCF motifs which had less than 20th percentile of CTCF binding signal. Among the kept CTCF motifs, we extended the CTCF motif interval by ± 10kb. For each jet, we then looked for intersecting CTCF motifs that lies in the neighborhood of a jet, where the neighborhood is defined by the jet extrusion point (**Figure S6g**). The CTCF motif(s) was categorized on its relative position with respect to the jet loading site (“upstream” or “downstream”) and whether the CTCF motif orientation is “forward” or “reverse”, yielding 4 possible categories: “upstream forward”, “upstream reverse”, “downstream forward”, and “downstream reverse”. We refer to a single jet and its intersecting CTCF motif(s) as a “cluster”. Notably, within a cluster we may have different combinations of categories (e.g., one jet may intersect with three CTCF motifs, one that is “upstream forward” and two that is “downstream reverse”). Suppose for simplicity’s sake we describe the procedure for a single “cluster”: we pooled together all the angles that align with the CTCF motif interval using the angle extraction scheme shown in **Figure S6g**, and computed the mean of the angles separately for each of the categories, plotting them as separate boxplots (**Figure 6j, Figure S6i**). As an example, consider a cluster that is a single jet intersecting with three CTCF motifs – one “upstream forward” and two “downstream reverse” – we compute the mean angles separately for each category (e.g., 84°, 96° respectively), then this jet contributes 84° to the “upstream forward” boxplot and 96° to the “downstream reverse” boxplot only once.

This way of pooling the angle points and taking the mean is repeated for each jet cluster and ensures that we do not inflate the differences between the boxplots as samples are independent within a category. We compare the boxplots by a two-sided Wilcoxon rank-sum test using `ranksums` from *scipy* (1.16.1). GM12878 STORM motifs (Tang et al.,, 2015) and CTCF ChIP-seq were used for both jets called in GM12878 (**Figure 6j**) and K562 Hi-C (**Figure S6i**) as CTCF boundaries are conserved across human cell-lines. The intersection and mean angle computation was done using *pandas* (2.2.2).

#### Jet angle deviation

The jet angle deviation is the mean absolute difference of jet angles from 90°. For the *k*th jet with length *z*_*k*_ and jet angles (**a**_*k*_(1), **a**_*k*_(2), …, **a**_*k*_(*z*_*k*_)), the jet angle deviation is

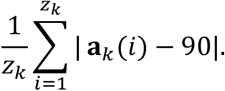

#### Defining GM12878 cohesin loading sites and anchor sites

As means of characterizing the jets identified by MIA-Jet with respect to cohesin loading and anchoring sites, 10239 cohesin-specific peaks from GM12878 cell line are further filtered by the criteria that NIPBL binding (of GSM2443453 downloaded track) is greater than 5 counts, yielding 5,467 regions. A total 3,014 convergent loops include 1 or more of these cohesin loading regions; since convergent loops may overlap (e.g., a bigger loop enclosing a smaller loop), some loading regions are recorded for two different convergent loops. These 3014 loops enclosed 8777 loading regions. We only kept loading sites that are within 25 percentile to 75 percentile from both left and right anchors, to exclude those near the anchors. The result is 2226 loops containing 4938 loading sites for GM12878 cells, but some loops have multiple loading sites and a loading site may be encapsulated inside multiple loops. As a final filter, we retained only 2352 unique loading sites and 3079 unique anchoring sites, which are used for overlapping MIA-Jet jets midpoint location extended both sides by 10 kb or random regions with *bedtools* `intersect -wa -u`.

#### Annotating GM12878 jets with chromatin states

MIA-Jet jets midpoint location extended both sides by 10 kb or random regions are initially overlapped against chromHMM states to obtain all states and basepairs of overlaps in each region via *bedtools* `intersect -wao`. The chromatin states 1,2,3 are grouped into ‘promoters’, 4,5,6,7 into ‘enhancers’, 9,10,11 into ‘transcribed states’, while 8 is ‘insulator’, 12 is ‘repressed’, 13 is ‘heterochromatin’, and the rest is classified as ‘other’. Given the number of basepairs occupied by each of these 6 simplified states, if the promoter is larger than 0 bp, then the state is ‘prom’. If the region does not contain any promoters and the enhancer is larger than 0 bp, then the state is ‘enh’; otherwise, the state is assigned to the one with maximum occupancy in terms of basepairs.

## DATA AND CODE AVAILABILITY

No experimental data have been generated in this study. Data used in this study are publicly available as stated in the Supplementary Table S1. The code (v2.0.3) and software is available under the MIT license on github: https://github.com/minjikimlab/miajet/tree/v2.0.3.

## ACKNOWLEDGEMENTS

This study was supported by the National Human Genome Research Institute (R00-HG011542). The authors thank Zhengrong Zhangding, Jiazhi Hu, Tony Lindeberg, Matthias Merkenschlager, and members of Minji Kim’s research group for helpful discussions; Laya Pullela and Zachary Apell for running Juicer on the DP thymocyte Hi-C data. We also thank Erez Lieberman-Aiden for depositing the intact Hi-C data to the ENCODE data portal for the public and sharing his thoughts on the cohesin extrusion model.

## AUTHOR CONTRIBUTIONS

M.K. conceptualized the project. S.K. devised algorithms, wrote the MIA-Jet Python package, and performed the benchmark with input from M.K. M.K. and S.K. interpreted results and wrote the manuscript.

## SUPPLEMENTARY FIGURE TITLES AND LEGENDS

**Figure S1:**
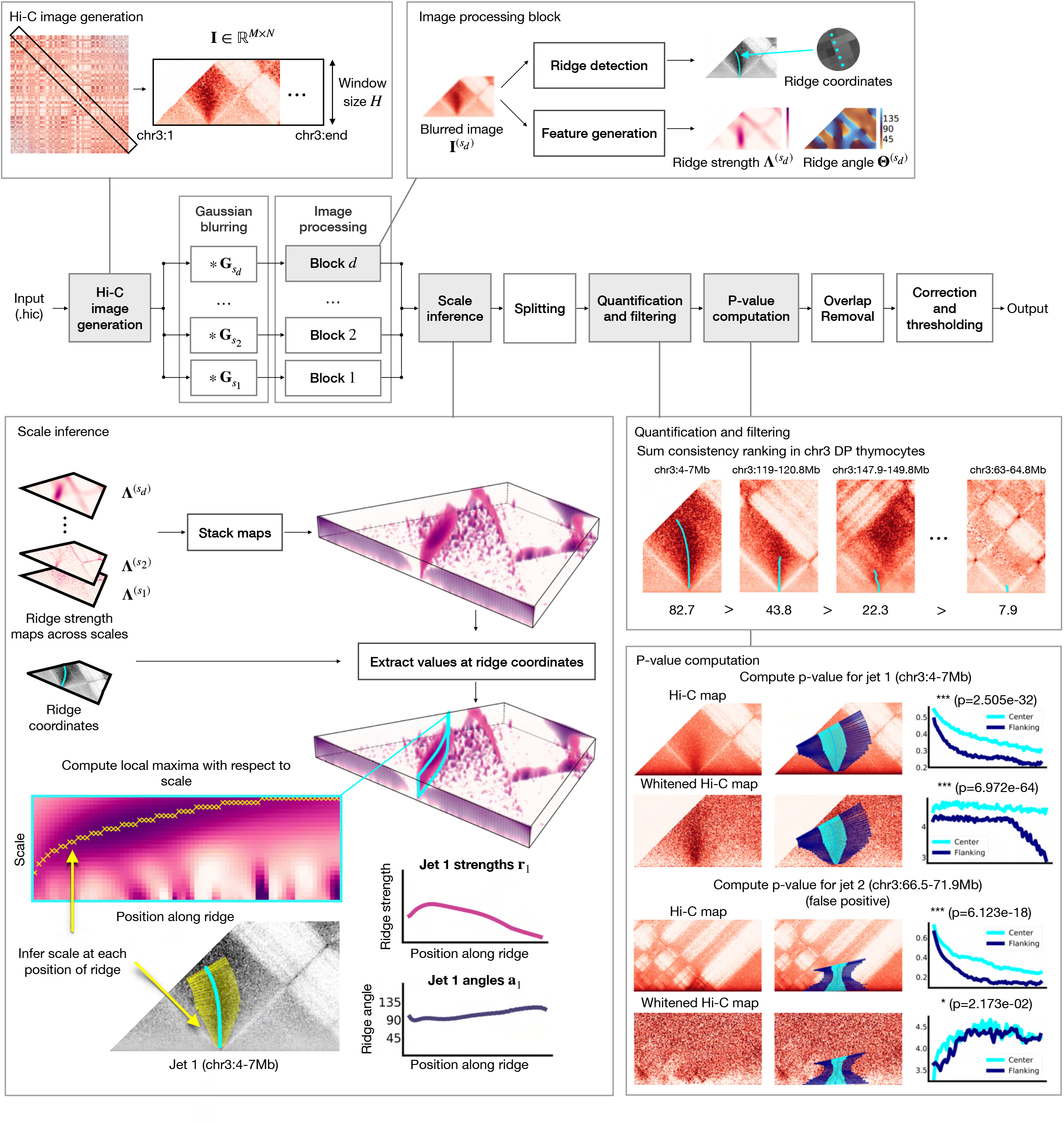
Detailed pipeline of MIA-Jet algorithm. Related to **Figure 1**. The MIA-Jet pipeline takes input a .hic contact map and extracts an upper-triangle rectangular matrix I ∈ ℝ^*M*×*N*^ (Hi-C image generation). The image is then fed into *d* blocks each applying a Gaussian blur with standard deviation *s*_1_ ≤ *s*_2_, ≤ *⋯* ≤ *s*_*d*_. The blurred image is passed into an image processing block which (1) applies ridge detection to infer ridge coordinates and (2) computes ridge strength and ridge angle from the blurred image (Image processing block). The ridge strength map is combined by stacking across different scales (Scale inference) to form a 3 dimensional tensor such that the z-axis represents each scale. We extract values at ridge coordinates to obtain a matrix that is scale × position along ridge, and infer the true scale at each position of the ridge (**Methods**, *Scale inference*). The inferred scales are used to compute the ridge strength signal **r**_*k*_ and angles **a**_*k*_, which are shown for example jet *k* = 1 (Scale inference). The jets are then processed and quantified, ranking jets by sum consistency metric (Quantification and filtering). P-values are computed on the Hi-C map comparing the values at the center region and flanking region (one-sided KS test), which is significant for jet 1 (P-value computation, row 1). On the whitened Hi-C map that reduces the strength of A/B compartments, jet 1 is still significant (P-value computation, row 2). For a false jet 2, the p-value computed on the Hi-C map is very significant due to the A/B compartment plaid pattern (P-value computation, row 3) but can be filtered out using the p-value computed on the whitened Hi-C map, which has larger p-value (P-value computation, row 4). Then, overlapping jets are resolved to call one jet, after which p-values are corrected and filtered. The output is a set of filtered jets with position, width, and angles, as well as other metrics (**Methods, Outputs**).

**Figure S2:**
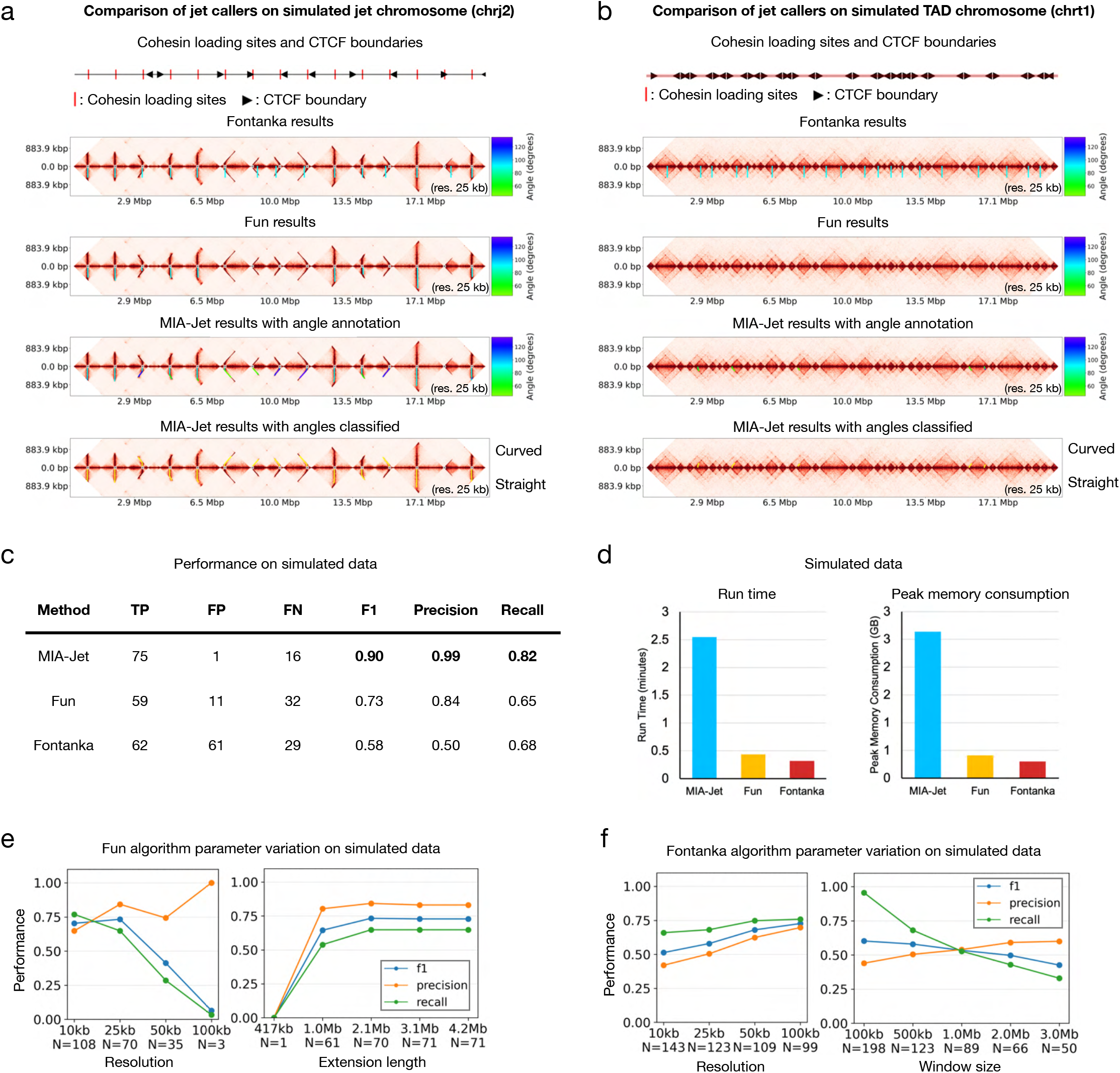
Examples of simulated data and parameter variation for Fun and Fontanka. Related to **Figure 2**. **(a)** Example of simulated chromosome chrj2 containing jets (‘test1cii’ **Methods**, *Generating simulated data*). Location of ground truth jet loading sites and CTCF boundary elements (row 1). Jets called by Fontanka (cyan line) (row 2), Fun (cyan line) (row 3), and MIA-Jet (coloured by angle) (row 4). Jets called by MIA-Jet classified into “Curved” (upper half) or “Straight” (lower half) using angle annotations (row 5). **(b)** Similar to panel **(a)** but for simulated chromosome chrt1 containing TADs (‘test1eii’ **Methods**, *Generating simulated data*). **(c)** Table showing performance of jet callers: True Positive (TP), False Positive (FP), False Negative (FN). **(d)** Run-time and peak memory consumption on simulated dataset. **(e)** Performance of Fun across parameter variation on simulated data. Extension length specifies maximum length of jets. **(f)** Performance of Fontanka across parameter variation on simulated data. Window size is the size of the jet mask kernel.

**Figure S3:**
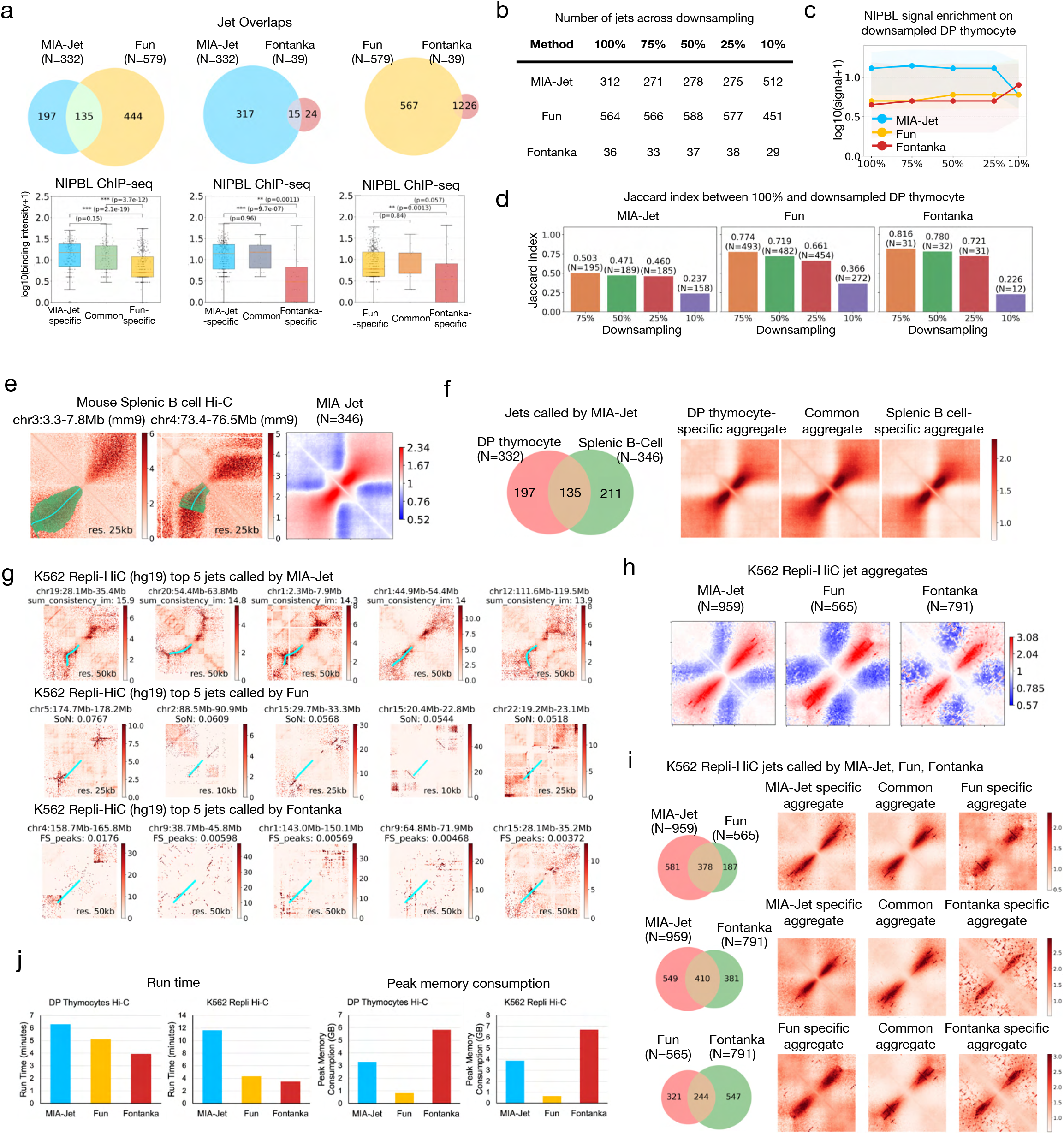
Examples and analyses of jets in DP thymocytes Hi-C, Splenic B cells Hi-C, and K562 Repli-HiC data. Related to **Figure 3**. **(a)** Comparison of jets called between MIA-Jet, Fun, and Fontanka. Venn diagram shows the number of jets common and specific between two methods. The NIPBL ChIP-seq boxplot is shown for the common and specific jets below. **(b)** Number of jets called by MIA-Jet, Fun, and Fontanka in downsampled DP thymocyte Hi-C data. **(c)** NIPBL ChIP-seq signal at jet locations called by jet callers in downsampled DP thymocyte Hi-C data. Shaded contour shows 25^th^ and 75^th^ percentiles. **(d)** Jaccard index between original (100%) DP thymocyte Hi-C and downsampled (75%, 50%, 25%, 10%) datasets for each jet caller. **(e)** Left and middle: Example jets called by MIA-Jet in splenic B cell with MIA-Jet annotation in lower triangle. Right: Aggregate Observed/Expected map of jets in splenic B cell. **(f)** Left: Venn diagram of jets between DP thymocytes and Splenic B cell Hi-C data. Right: Aggregate Observed/Expected maps at each of the three categories of jet regions. **(g)** Top 5 jets identified by MIA-Jet (row 1), Fun (row 2), Fontanka (row 3) in K562 Repli-HiC. Maps show Observed/Expected values and annotation in lower triangle. **(h)** Aggregate Observed/Expected maps of jets identified by MIA-Jet, Fun, and Fontanka in K562 Repli-HiC. **(i)** Comparison of jets identified by MIA-Jet, Fun, and Fontanka, in K562 Repli-HiC data with Venn diagram on the left and 2D contact aggregation on the right. **(j)** Run-time and peak memory consumption on DP thymocyte Hi-C and K562 Repli-HiC data.

**Figure S4:**
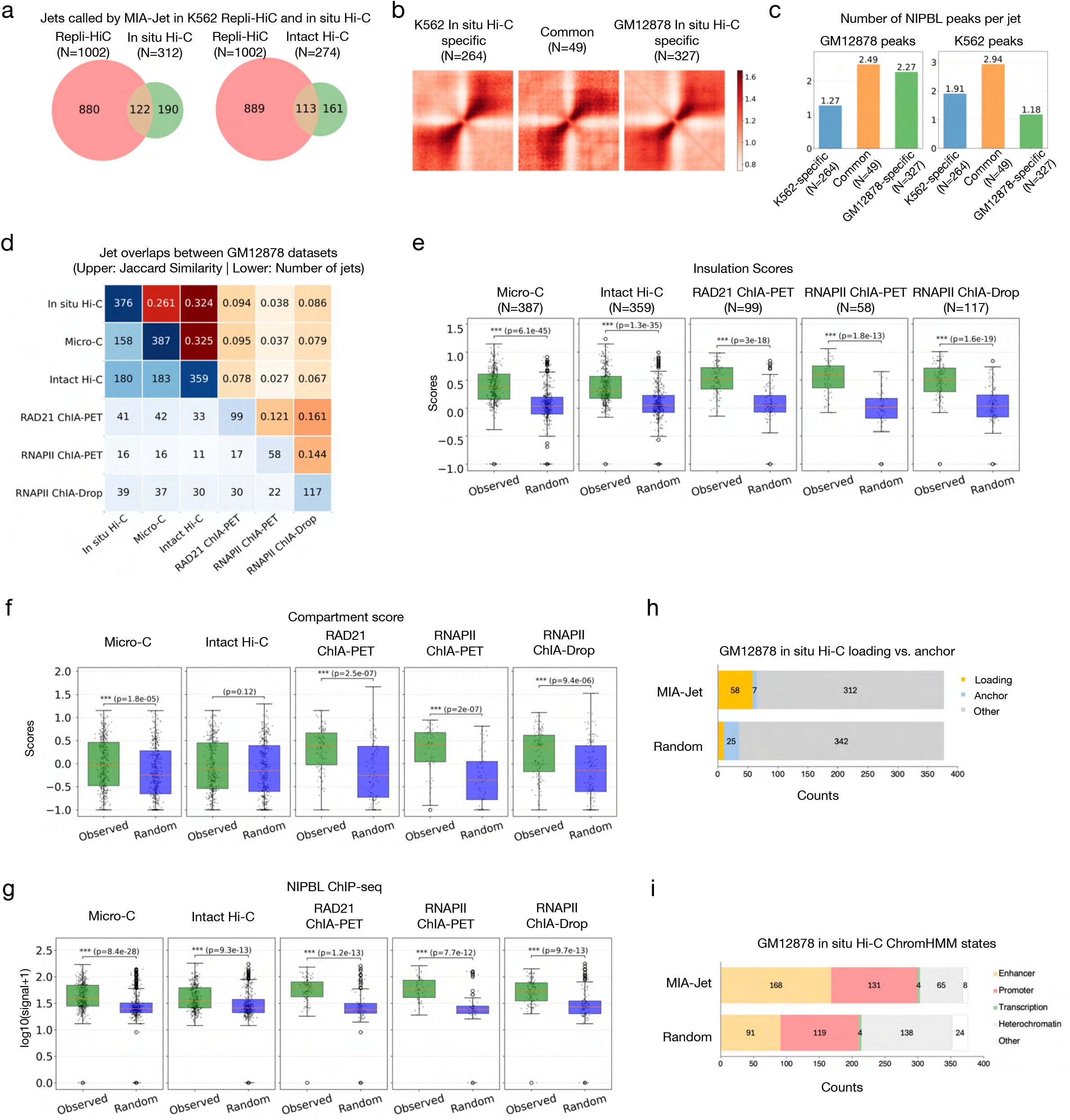
Characterization of jets identified by MIA-Jet in K562 and GM12878 cell lines. Related to **Figure 4**. **(a)** Venn diagrams comparing jets identified between K562 Repli-HiC and in situ Hi-C (left) or intact Hi-C (right). **(b)** Aggregate Observed/Expected maps at common and specific jet regions comparing K562 in situ Hi-C and GM12878 in situ Hi-C; N is the number of locations in each category. **(c)** Number of NIPBL peaks at jet loading site at common and specific jets comparing K562 in situ Hi-C and GM12878 in situ Hi-C. Left: GM12878 NIPBL peaks. Right: K562 NIPBL peaks. **(d)** Pairwise comparison matrix of jet overlaps among GM12878 datasets. Lower triangle: Number of jets overlapping between two datasets. Upper triangle: Jaccard index. Diagonal: Number of jets called in each dataset. **(e)** Boxplots of insulation scores at jet locations (Observed) and random locations (Random) across GM12878 technologies (p-values from two-sided Wilcoxon rank-sum test). **(f)** Boxplot of compartment scores at same locations as panel **(e). (g)** Boxplots of log-transformed NIPBL ChIP-seq data at same locations as panel **(e). (h)** Stacked bar chart of jets associated with cohesin loading (yellow), CTCF anchoring (blue), or other regions for jet locations (MIA-Jet) and random locations. **(i)** Composition of chromHMM chromatin states at jet origins (see **Methods**, *Annotating GM12878 jets with chromatin states*).

**Figure S5:**
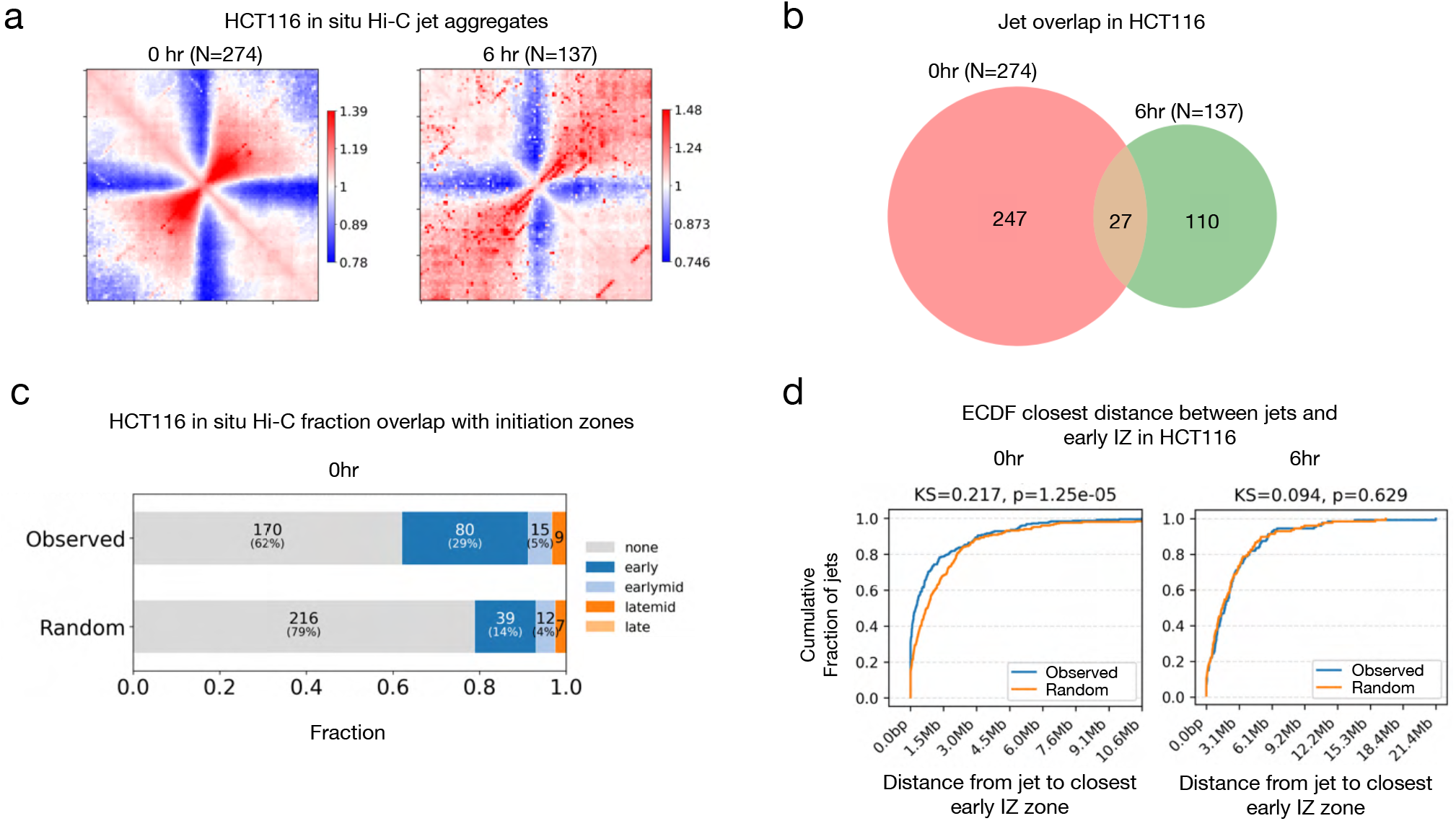
Analysis of jets identified in cohesin depleted cells. Related to **Figure 5**. **(a)** Aggregate Observed/Expected map of jets called in HCT116 0hr (left) and 6hr (right). ‘0h’: without auxin treatment (before depletion; control), ‘6h’: 6 hours of auxin treatment (RAD21 depletion). **(b)**. Venn diagram of jets called in HCT116 0hr and 6hr. **(c)** Stacked bar chart of jets associated with replication initiation zone annotations for jet locations called by MIA-Jet (Observed) and random locations. **(d)** Empirical cumulative distribution function of distance from “Observed” or “Random” locations to closest early initiation zones in HCT116 0hr (left) 6hr (right). Observed: jets locations identified by MIA-Jet. Random: random regions. P-values from two-sided Kolmogorov-Smirnov test.

**Figure S6:**
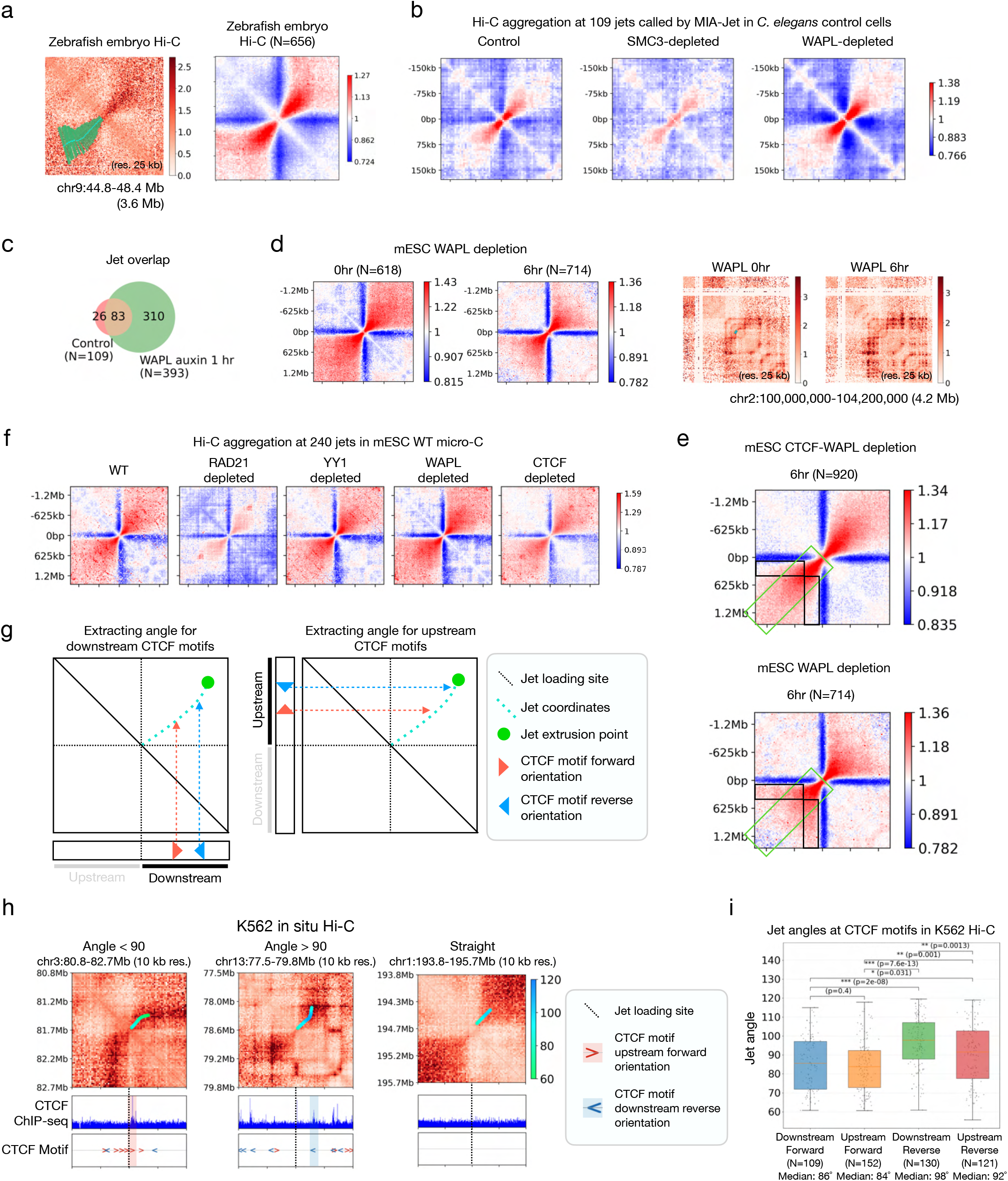
Analysis of protein depletions on jet extrusion. Related to **Figure 6**. **(a)** Left: Example jet identified by MIA-Jet in zebrafish embryo Hi-C data. Right: Aggregate Observed/Expected map of jets called in zebrafish embryos. **(b)** Aggregate Observed/Expected maps at 109 jet regions identified in *C. elegans* control Hi-C data, shown also for SMC3-depleted, and WAPL-depleted cells. **(c)** Venn diagram of jets called in *C. elegans* control and WAPL-depleted cells. **(d)** Left: Aggregate Observed/Expected maps of jets called in mESC WAPL 0hr (control) and 6hr (WAPL depletion). Right: Example region. **(e)** Same data as panel **Figure 6c** and panel **(d)** but highlighting the differences in enrichment between CTCF-WAPL DKO and WAPL KO. Enrichment in CTCF-WAPL DKO relative to WAPL KO is highlighted in a green box and depletion in a black box. **(f)** Aggregate Observed/Expected maps at 240 jet regions identified in WT mESC Micro-C data, shown also for protein depleted cells: RAD21, YY1, WAPL, and CTCF depleted. **(g)** Schematic depicting how to find jet points that intersect with CTCF motifs. Left: CTCF motifs downstream of jet loading site. Right: CTCF motifs upstream of jet loading site. Each jet point is associated with an angle, annotated from the MIA-jet program. **(h)** Example jets with MIA-Jet line annotation coloured by angles in K562 in situ Hi-C data. Left: jet angled less than 90° with corresponding CTCF motif upstream and forward orientation in red highlight. Middle: jet angled greater than 90° with corresponding CTCF motif downstream and reverse orientation in blue highlight. Right: straight jet with no intersecting CTCF motifs. **(i)** Boxplots of mean jet angle corresponding to CTCF motifs (downstream forward, upstream forward, downstream reverse, and upstream reverse) for K562 Hi-C data. N is the number of jets intersecting with each CTCF motif category. P-values from two-sided Wilcoxon rank-sum test.

